# AreTomoLive: Automated reconstruction of comprehensively-corrected and denoised cryo-electron tomograms in real-time and at high throughput

**DOI:** 10.1101/2025.03.11.642690

**Authors:** Ariana Peck, Yue Yu, Mohammadreza Paraan, Dari Kimanius, Utz H. Ermel, Joshua Hutchings, Daniel Serwas, Hannah Siems, Norbert S. Hill, Mallak Ali, Julia Peukes, Garrett A. Greenan, Shu-Hsien Sheu, Elizabeth A. Montabana, Bridget Carragher, Clinton S. Potter, David A. Agard, Shawn Zheng

## Abstract

A high throughput processing pipeline that performs comprehensive corrections is needed to realize the full potential of cryo-electron tomography and subtomogram averaging. The field’s fragmented software landscape remains a significant hurdle to this end. Here we present AreTomoLive, an automated real-time pipeline composed of two GPU-accelerated packages. The first, AreTomo3, streamlines tomographic alignment and reconstruction, with new features to fully account for sample geometry, locally correct the contrast transfer function, and curate data for downstream tasks. The second package, DenoisET, is a new implementation of the machine learning algorithm Noise2Noise and runs in parallel with AreTomo3 to perform contrast enhancement. To reduce barriers to routine use, AreTomoLive prioritizes automation: AreTomo3 autonomously pauses and reactivates processing depending on the status of data collection, while DenoisET algorithmically determines when to transition from training to inference. AreTomoLive endeavors to advance cryoET for *in situ* structural analysis with its comprehensive corrections and full automation.

## Introduction

Cryo-electron tomography (cryoET) is a powerful technique for visualizing the molecular machinery that drives cellular function. Recent studies have highlighted the ability of this method to resolve molecular structures with up to sub-nanometer resolution, importantly while preserving detailed cellular context^1–3^. However, a survey of the Electron Microscopy Data Bank^4^ shows that sub-nanometer resolution has only been achieved a handful of times for molecules *in situ*, and primarily for abundant or high-symmetry species (Extended Data Fig. 1). Much of the difficulty stems from the challenge of scale: in particular, the need to identify tens of thousands of copies of the target molecule in a crowded cellular environment across tens to many hundreds of tomograms, which are 3D maps of the imaged sample. Once located, these copies must be accurately aligned and averaged in a process called subtomogram averaging (STA) to determine the molecule’s structure in its native cellular environment^5^.

Improvements in throughput, alignment accuracy, and contrast are critical to extend STA to new molecular species. Higher throughput is needed to locate sufficient copies of the target molecule for STA to overcome the extremely low signal-to-noise ratio (SNR) of individual and often conformationally heterogeneous molecules. Parallel data acquisition schemes like PACE-Tomo have enabled the routine collection of more than 1000 tilt-series per week on a single electron microscope^6–9^, but realizing the full potential of this accelerated acquisition requires data processing to keep pace. Alignment accuracy is a second key consideration given that biological samples deform upon exposure to an electron beam^10^, and the signal from the sample will be further delocalized by the microscope’s contrast transfer function (CTF)^11,12^. Left uncorrected, these aberrations and beam-induced motion (BIM) severely limit the high-resolution information contained in cryoET data and impede high-resolution STA. Finally, contrast enhancement during data processing is often critical to locate target molecules both by manual annotation and automated picking algorithms and to interpret their cellular context^13^.

The software landscape to perform cryoET data preprocessing — spanning motion correction of the acquired tilt-movies to contrast enhancement of the reconstructed tomograms — remains fragmented, making it difficult to achieve high-throughput. Packages such as M^1^, Scipion^14^, and TomoBEAR^15^ have been developed to provide an end-to-end workflow. However, these pipelines largely assemble third-party software packages that rely on distinct coordinate conventions, file formats, and software dependencies, making them difficult to run and maintain. Further, intermediate data from each package must be saved to and re-read from disk, creating unwanted overhead for processing at scale. To address these barriers, AreTomoLive provides a single integrated pipeline that automates each step of data preprocessing and introduces several innovations to improve alignment and reconstruction accuracy. The first component of this pipeline, AreTomo3, streamlines the major functions of the widely-used software packages MotionCor2^16^ and AreTomo^17^ to correct for global and local BIM in both 2D and 3D in addition to integrating new features to robustly determine the defocus handedness, measure sample geometry, locally correct the CTF, and curate data. The second component, DenoisET, is a new implementation of the machine learning (ML) algorithm Noise2Noise^18^ that leverages the comprehensively-corrected tomograms generated by AreTomo3 to provide on-the-fly 3D denoising.

The design of AreTomoLive prioritizes throughput, automation, and versatility. In terms of throughput, AreTomoLive can catch up with tomographic data collection configured for 12 targets per stage position when run on a Linux server equipped with 8 NVIDIA RTX A6000 GPUs. To achieve automation, AreTomo3 pauses and reactivates processing as the status of data collection changes, while DenoisET algorithmically determines both when sufficient high-quality tomograms are available for training and when to transition from training to inference. For versatility, AreTomoLive provides multiple entry points to facilitate data reprocessing and generates processed data files and metadata in different formats depending on the downstream task. We hope that this pipeline will meet the field’s pressing need for a high-throughput preprocessing pipeline to realize the full potential of cryoET for *in situ* structural analysis at scale.

## Results

### Real-time and autonomous tomographic alignment and reconstruction

AreTomo3 offers a streamlined workflow to reconstruct tomograms from raw tilt-movies and is divided into three modules (Fig. 1, dark blue arrows). The first module inherits corrections for anisotropic BIM in 2D from Motion-Cor^16^. Motion-corrected micrographs are then sorted by tilt angle to assemble the tilt-series. The second module performs iterative CTF estimation and tomographic alignment, extending the alignment procedure developed in AreTomo^17^. This module first estimates the per-tilt CTF parameters before performing global tomographic alignment, which defines the tilt axis angle and corrects for the relative translations between the tilt images. The tilt axis angle is then taken into account to refine the CTF parameters and determine the pre-tilt angular offsets *α*_0_ and *β*_0_, which collectively give rise to an oblique defocus gradient across each tilted image (Extended Data Fig. 2). This module next measures the sample thickness before computing the local alignments to correct for 3D deformations that occur in subregions across the full tomographic volume. The third and final module applies the global and local alignments to the tilt-series to reconstruct tomograms either by the simultaneous algebraic reconstruction technique (SART)^19^ or weighted backprojection^20^, with corrections for the local CTF and *α*_0_ offset optionally applied. In addition to tomograms, AreTomo3 generates additional files to facilitate downstream processing. These include a record of the alignments applied to the tilt-series during reconstruction, the tilt-series and per-tilt CTF parameters required by STA workflows like RELION-5^21^, and quality metrics that characterize each processed tilt-series.

**Figure 1:**
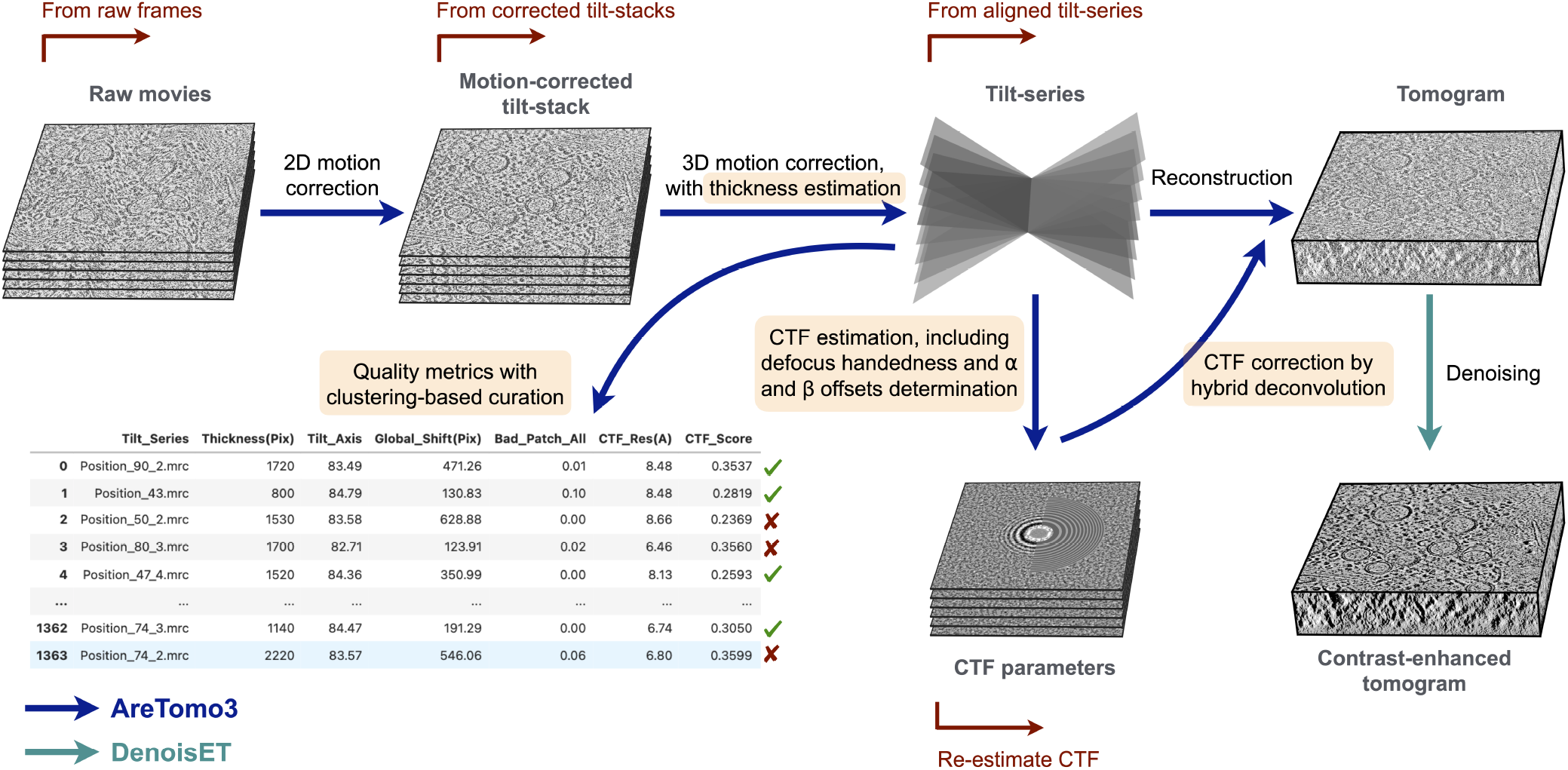
AreTomoLive provides a fully automated pipeline for tomographic alignment, reconstruction, and contrast enhancement in real time. AreTomoLive is composed of two GPU-accelerated packages, AreTomo3 and DenoisET. AreTomo3 performs the tasks indicated by the dark blue arrows to provide comprehensively-corrected tomograms minutes after each raw tilt-movie is collected. Beyond integrating the MotionCor2^16^ and AreTomo^17^ software packages, AreTomo3 introduces new features (highlighted text) to improve the accuracy of alignment and reconstruction and facilitate data curation. The red arrows indicate the multiple entry points that allow users to efficiently reprocess data from an intermediate state. The second component of AreTomoLive, DenoisET, runs in parallel with AreTomo3 to train and then apply an ML-based denoising model to the tomograms that AreTomo3 continually writes to disk (light blue arrow).

Parallel acquisition schemes^8,9^ have significantly increased the throughput of data collection and raised the bar for data preprocessing to provide users with feedback in real time — i.e., concurrent with data collection. In addition to speed, flexibility is desired to let users pause acquisition to queue more targets and change collection settings. AreTomo3 achieves these design objectives through two principle mechanisms. First, delivering an integrated pipeline rather than stitching packages together allows AreTomo3 to retain data in system memory, reducing the number of times large data files have to be read from disk. Second, AreTomo3 implements a scout-worker system like Warp’s^22^ in which the scout thread assigns any new raw tilt-series it detects to idle worker threads, which process data independently and concurrently. This design allows AreTomo3 both to scale with the number of allocated GPUs (Table 1) and to run autonomously, entering a latent state if processing outpaces data collection and reactivating worker threads as new tilt-series are found. Extended Data Fig. 3 tracks AreTomoLive’s performance during a multi-day data collection session, showing that AreTomo3 and DenoisET completed processing all collected tilt-series within minutes of the session’s end. In addition to real-time processing, AreTomoLive performs offline processing and offers multiple entry points that allow users to bypass upstream tasks when reprocessing data with adjusted alignment or reconstruction parameters (Fig. 1, red arrows).

**Table 1:**
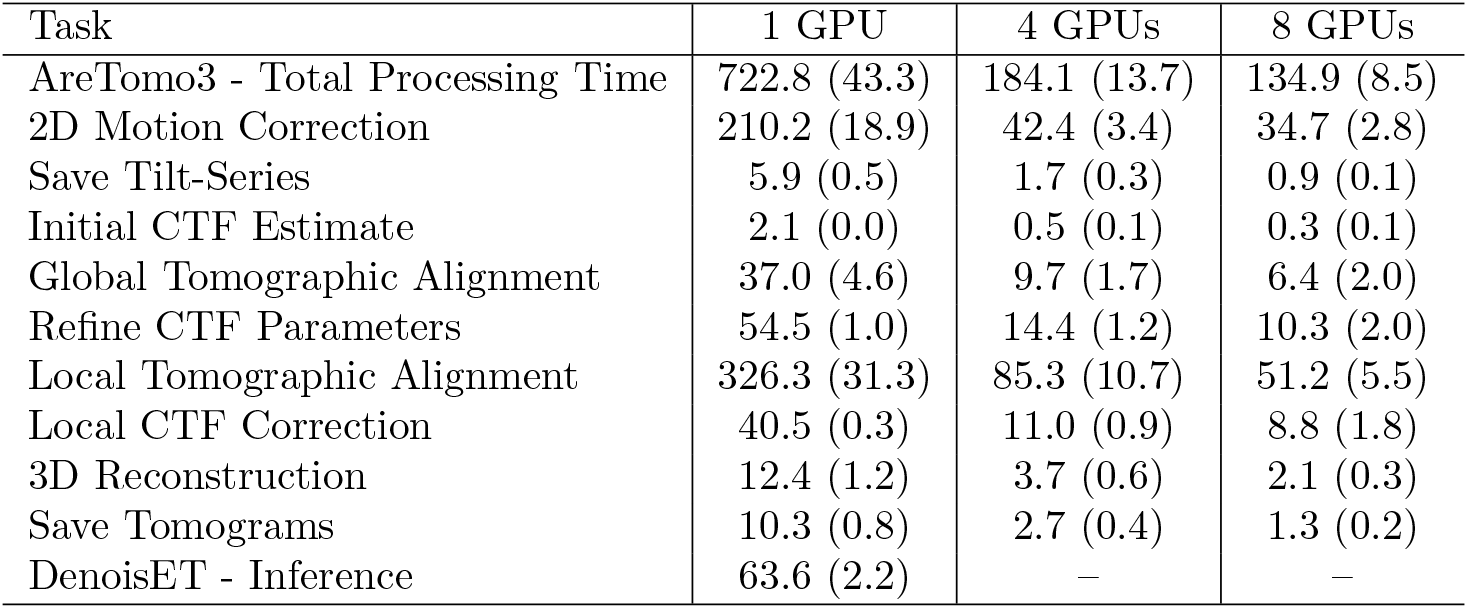
AreTomoLive’s performance and scalability. Performance was measured on the indicated number of NVIDIA RTX A6000 GPUs. The time required to complete each processing task per tilt-series is normalized by the number of GPUs and listed in seconds, with the standard deviation from processing *>*100 tilt-series noted in parenthesis. The raw data were collected at a pixel size of 1.54 Å. 2D motion correction with 4 × 4 local patches was performed on super-resolution EER movies, followed by downsampling to the original pixel size before performing tomographic alignment with 4 × 4 patches. CTF-corrected even, odd, and full tomograms were reconstructed at a pixel size of 5 Å and a volume depth of 250 nm. Near-linear scaling was observed on 4 GPUs, with a slight reduction on 8 GPUs due to increased overhead. Currently, DenoisET does not support parallelization.

| Task | 1 GPU | 4 GPUs | 8 GPUs |
| --- | --- | --- | --- |
| AreTomo3 - Total Processing Time | 722.8 (43.3) | 184.1 (13.7) | 134.9 (8.5) |
| 2D Motion Correction | 210.2 (18.9) | 42.4 (3.4) | 34.7 (2.8) |
| Save Tilt-Series | 5.9 (0.5) | 1.7 (0.3) | 0.9 (0.1) |
| Initial CTF Estimate | 2.1 (0.0) | 0.5 (0.1) | 0.3 (0.1) |
| Global Tomographic Alignment | 37.0 (4.6) | 9.7 (1.7) | 6.4 (2.0) |
| Refine CTF Parameters | 54.5 (1.0) | 14.4 (1.2) | 10.3 (2.0) |
| Local Tomographic Alignment | 326.3 (31.3) | 85.3 (10.7) | 51.2 (5.5) |
| Local CTF Correction | 40.5 (0.3) | 11.0 (0.9) | 8.8 (1.8) |
| 3D Reconstruction | 12.4 (1.2) | 3.7 (0.6) | 2.1 (0.3) |
| Save Tomograms | 10.3 (0.8) | 2.7 (0.4) | 1.3 (0.2) |
| DenoisET - Inference | 63.6 (2.2) | — | — |

### Tilt-series CTF estimation

Tilt-series data pose several challenges to robustly measuring the CTF parameters. First, though comparable to the dose used to acquire a single-particle cryoEM micrograph, the total dose per tilt-series is fractionated across the full tilt range, resulting in weaker Thon rings in individual tilt images. Second, the defocus fluctuates across the tilt-series due to imperfect tracking and auto-focusing. As a result, this parameter must be refined for each tilt image. Third, the increased effective sample thickness at high tilt angles attenuates the Thon rings. This diminished signal is compounded by the oblique defocus gradient that arises from the combination of the tilt angle and the *α*_0_ and *β*_0_ offsets and reduces the spatial coherence of the image’s power spectrum (Extended Data Figs. 2 and 4).

To address these challenges, AreTomo3 introduces several new innovations to adapt traditional approaches to CTF fitting for tilt-series data. One of these new features is the self-adaptive determination of the defocus search range and step size since choosing values that achieve a good balance between accuracy and speed can be difficult. Further, the optimal values are related to the magnification, which spans a broader range in cryoET than single particle cryoEM data collection. AreTomo3 automates the selection of these parameters based on the magnification change given by:

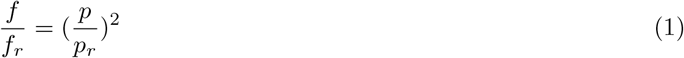

where *f, p*, and the subscript *r* denote the defocus, pixel size, and reference, respectively (see Methods). Based on a default search range and step size at a reference magnification that yields a pixel size of 1 Å, AreTomo3 rescales these parameters by the square of the corresponding pixel size to accommodate different magnifications.

A second innovation is the joint optimization of the tilt offsets, tilt axis ambiguity factor (see next section), and per-tilt CTF parameters. AreTomo3 achieves this joint optimization by dividing each tilt image into overlapping tiles and computing the power spectrum of each tile. For each tilt image *i*, the defocus at each tile center *f*_*ij*_ will experience a shift relative to the defocus at the image center *f*_*i*_ according to:

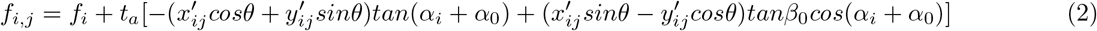

where *α*_*i*_ denotes the nominal tilt angle, *θ* is the tilt axis angle, *α*_0_ and *β*_0_ are the sample’s tilt offsets at nominal 0° (Extended Data Fig. 2), 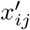 and 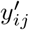 are the tile’s center coordinates, and *t*_*a*_ is the tilt axis ambiguity factor.

Following the approach in BSoft^23^, the power spectrum of each tile is rescaled based on its local defocus:

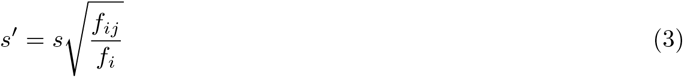

where *s* and *s* ^′^ are the spatial frequencies before and after rescaling, respectively. By compensating for the defocus gradient, this spectral rescaling increases the spatial coherence of the tilt image’s average power spectrum. Are-Tomo3 then jointly optimizes the CTF parameters with the defocus constrained by *α*_0_, *β*_0_, and *t*_*α*_ to maximize the objective function^24^:

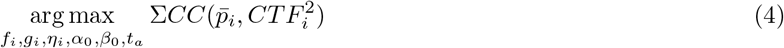

where 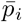 is the average of the rescaled power spectra of tiles extracted from image *i, CTF*_*i*_ is the predicted contrast transfer function for the tilt image, *CC* is the cross-correlation coefficient, and *g*_*i*_ and *η*_*j*_ are respectively the magnitude and azimuth angle of astigmatism. Because the rescaling factor in Eq. 3 depends on *f*_*i*_, optimization initially estimates this parameter before accounting for the tilting-induced defocus gradient or performing spectral rescaling. The optimal parameter set is found by a multiscale grid search followed by coordinate ascent.

A third adaptation adjusts the above approach to compensate for the attenuated Thon ring signal at higher tilt angles. Specifically, AreTomo3 bootstraps the astigmatism and, if a phase plate is used, additional phase shift parameters at tilt angles beyond ±30° to the values determined for the nearest less tilted image. Beyond this 30° threshold, only the defocus is estimated since this is expected to fluctuate more significantly between tilt images than the other parameters. Empirically we have observed that reserving the weaker signal at high tilt images to only refine the defocus improves convergence during optimization.

Collectively these adaptations have enabled robust CTF estimation for tilt-series data, including in challenging cases like data collected using a modified TYGRESS scheme^25^. In this scheme, the untilted image is not only collected at higher dose as in conventional TYGRESS but also close to focus to facilitate 2D template matching^26^, while the tilted images are acquired at higher defocus and low dose to balance the competing needs of increasing contrast for alignment and minimizing radiation damage. AreTomo3 was able to accurately fit the CTF for this dataset despite the dose discontinuity and large variation in defocus across the tilt-series (Extended Data Fig. 5).

### Determination of tilt axis directionality and defocus handedness

One drawback of the fragmented software ecosystem in cryoET is the use of different coordinate conventions, which can flip the defocus handedness when data are transferred between packages. Programs such as RELION-5^21^ and Warp^22^ define positive defocus handedness as the coordinate system in which an increased specimen tilt angle reduces the defocus of a particle with a positive *x* coordinate (Extended Data Fig. 6a). In this convention, the *z*-axis points toward the electron source. On the other hand, in a coordinate system with negative defocus handedness, increasing the tilt angle increases the defocus of a particle with a positive *x* coordinate. In this case, the *z*-axis points away from the electron source (Extended Data Fig. 6b). Switching between these coordinate systems flips the directionality of the sample rotation axis, resulting in a 180° error in the tilt axis angle. It is also possible for the directionality of the tilt axis to be incorrectly assigned during global alignment, which uses the common line approach that only determines the orientation of this axis but not its directionality^17^.

AreTomo3 adopts the coordinate system with a positive defocus handedness and resolves the tilt axis ambiguity factor, *t*_*a*_, with respect to this reference frame. Specifically, optimization of the CTF parameters compares *t*_*a*_=±1 given the initial estimate of the tilt axis angle (Eq. 2). The correct choice enhances the tiles’ spatial coherence during spectral rescaling and in turn yields a higher cross-correlation between the average power spectrum and the modeled CTF, an effect that will be most pronounced for tilt images with a large defocus gradient (Fig. 2). If *t*_*a*_ is determined to be negative, the tilt axis angle estimated by the common line approach is corrected by subtracting 180° to revert to a coordinate system with positive defocus handedness. AreTomo3 records the defocus handedness for each tilt-series it processes to ensure that downstream tasks can correctly interpret the imaging data and their corresponding alignments.

**Figure 2:**
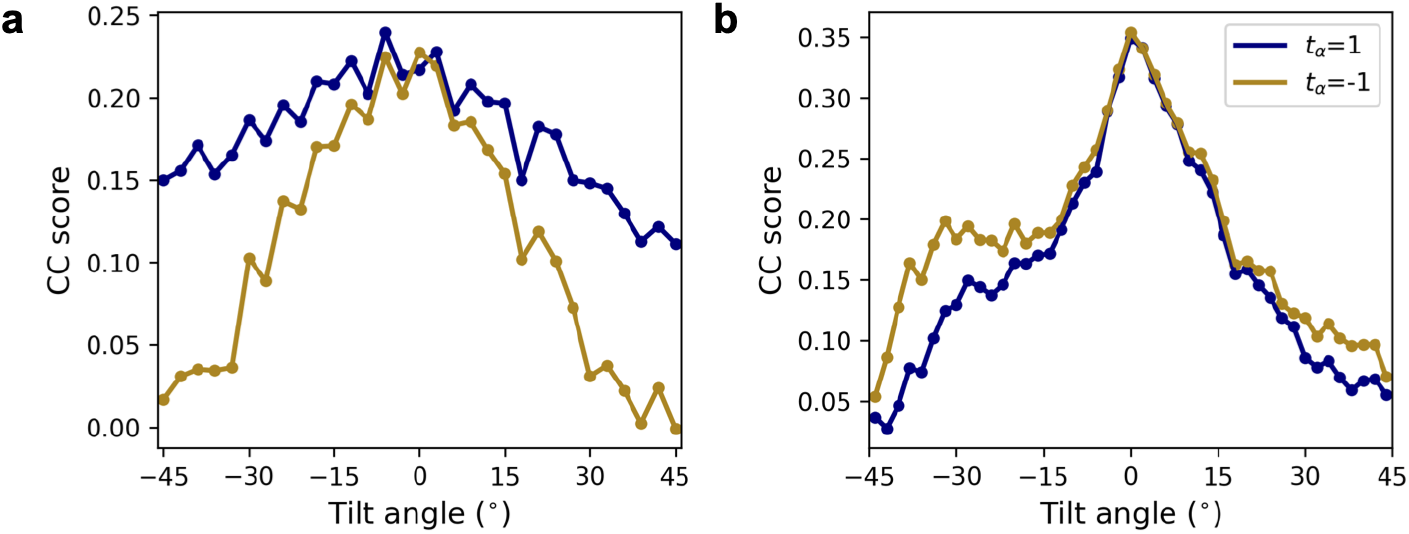
Resolution of the 180° ambiguity in the tilt axis angle from locally fitting the CTF. The sign of the tilt axis ambiguity factor, *t*_*a*_, was determined by comparing CTF scores, in particular for tilt images with a large defocus gradient. **a**. The tilt axis ambiguity factor was found to be positive, confirming the initial estimate of the tilt axis angle. **b**. The tilt axis ambiguity factor was determined to be negative, requiring correction of the tilt axis angle by subtracting 180°.

### Sample thickness estimation

AreTomo3 automates the measurement of sample thickness for more robust tomographic alignment. Previously in AreTomo^17^, users could specify a thickness value to be used during the alignment process. However, sample thickness is difficult to gauge prior to reconstruction, and a single value is generally unsuitable for batch processing due to the high variability in thickness among targets, including from the same sample. To measure sample thickness, AreTomo3 first reconstructs 10x binned tomograms by SART and then computes the cross-correlation between adjacent slices along the tomogram’s depth. These correlation profiles tend to have characteristic shapes, with higher values in the central region due to the continuity of structural information between adjacent *xy* slices within the sample (Fig. 3). The correlation decays to minima beyond the sample boundaries before increasing again near the tomogram edges due to backprojection artifacts. AreTomo3 determines the sample boundaries to be positioned where the correlation falls to the value halfway between the local maximum nearest the center of the profile and the minima on either side.

**Figure 3:**
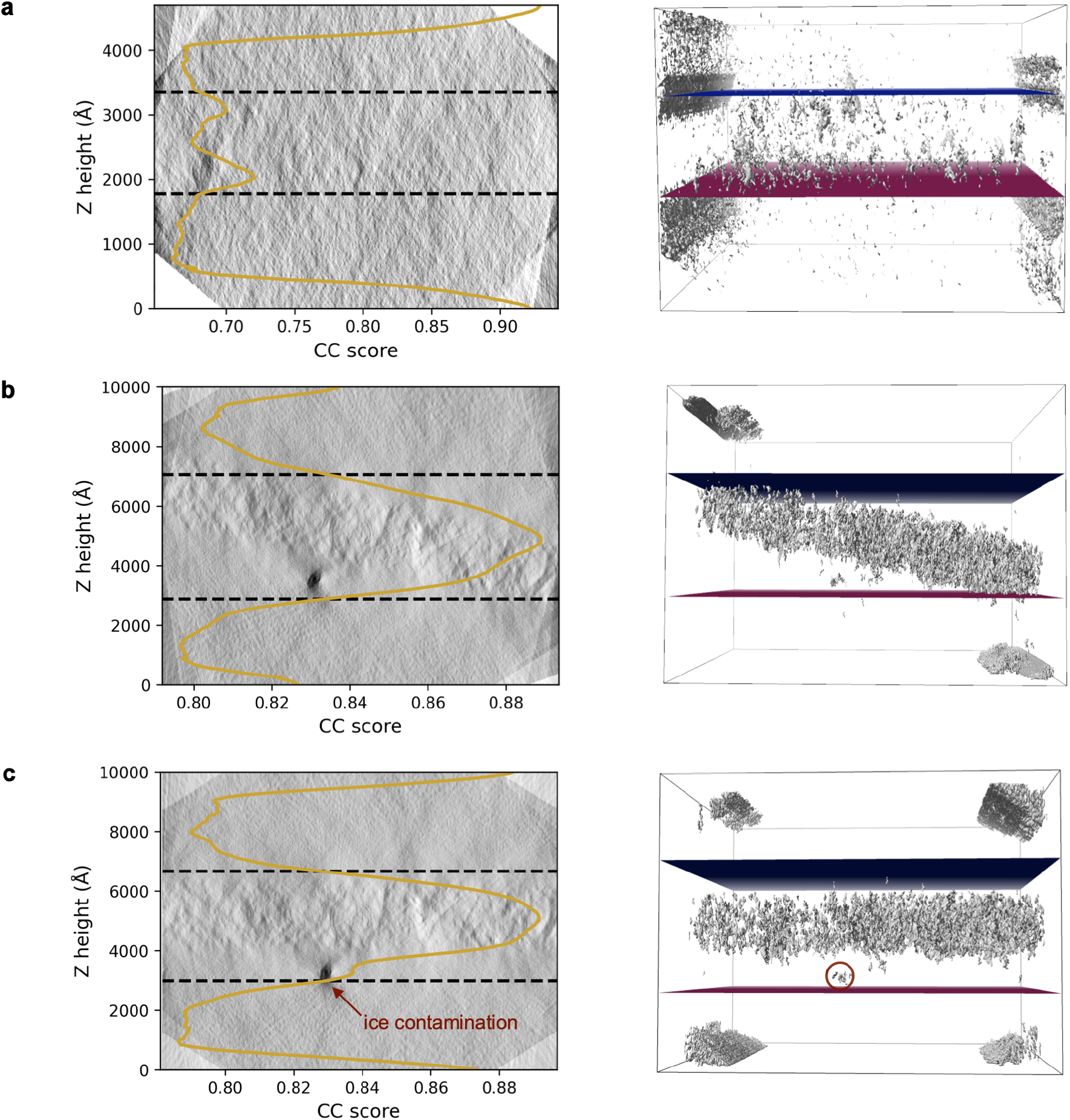
Runtime measurement of sample thickness. Slices through the *xz*-plane (left) and 3d views (right) are visualized for tomograms of **a**. a mixture of purified virus-like-particles and apo-ferritin and **b-c**. coronavirus-induced double-membrane vesicles. The cross-correlation (CC) scores between adjacent slices along the *z*-axis are overlaid in yellow and were used to estimate the sample boundaries (dashed lines and blue and magenta planes in the left and right panels, respectively). Correction of the *α*_0_ tilt observed in **b** levels the sample within the tomogram and improves the accuracy of the thickness measurement, as shown in **c**. However, the presence of a large ice crystal (red) artificially inflates the measured thickness.

This approach generalizes well to diverse samples, with the detected specimen boundaries fully encompassing the sample while excluding vacuum on either side (Fig. 3). However, this approach assumes that the sample is leveled within the tomogram. Consequently, samples like lamella that are characterized by a large *α*_0_ tilt should be reconstructed with this offset corrected to exclude vacuum above and below the inclined plane of the sample (Fig. 3b-c). AreTomo3 automatically accounts for the *α*_0_ tilt offset during alignment and can optionally do so during reconstruction as well. The boundary detection process can also be confounded by ice contamination, which can yield high cross-correlation scores even in regions with minimal structural information from the sample (Fig. 3c).

Correctly determining the sample thickness is critical for alignment accuracy. AreTomo3 computes local alignments by performing projection matching between the globally aligned tilt-series and a reference tilt-series that is generated by forward-projecting an intermediate tomogram. An overestimate of the sample thickness introduces vacuum into this intermediate tomogram, which in turn propagates noise into the reference tilt-series. Though generally less detrimental, an underestimate restricts the structural information available for the alignment process. Either scenario produces a poor reference tilt-series, consequently reducing the accuracy of the local alignments and deteriorating the quality of the final tomogram. As an example, aligning a tilt-series of purified synaptosomes using a sample thickness of roughly half or twice the true value yielded tomograms in which the synaptic vesicles and membranes were obviously misaligned and poorly resolved (Extended Data Fig. 7).

### Local CTF correction

Accurately correcting the CTF requires accounting for the defocus gradient. While 2D strip-based and 3D CTF corrections have been implemented^27–29^, the most rigorous approach is a 2D tile-based method that fully compensates for the oblique defocus gradient induced by the tilt angle and the *α*_0_ and *β*_0_ offsets prior to reconstruction. AreTomo3 performs this local correction by dividing each raw tilt image into a series of overlapping tiles, with apodization applied to avoid checkerboard artifacts. The CTF deconvolution described below is then performed on each apodized tile, with all tiles from each tilt image sharing the same per-tilt CTF parameters except for the defocus, which is calculated based on Eq. 2 to account for its local variation across the tilt image. The CTF-corrected tilt image is then reconstructed by stitching the central squares of adjacent tiles. Analogous to the approach taken in TomoAlign^30^, this strategy of deconvolving the CTF from a region larger than the one used during image reconstruction reduces signal delocalization and CTF aliasing in the high-frequency domain^31^.

CTF corrections can be broadly categorized into two approaches: phase flipping and Wiener filtering^32^. The former flips the sign of the Fourier components at the frequencies where the CTF is negative, consequently restoring the correct phases while preserving the original noise statistics. However, phase flipping does not compensate for CTF-induced amplitude oscillations, which can be significant in the low to medium frequency range. By contrast, Wiener filtering aims to correct both the amplitude and phase terms but requires estimating the spectral SNR (SSNR), which is difficult to do accurately. This approach also distorts the noise statistics, especially near the zero-crossings where a poorly estimated SSNR will dominate the filter. To balance these concerns, AreTomo3 implements a hybrid correction scheme that transitions from a modified Wiener filter at low spatial frequencies to phase flipping at high frequencies, where the filter that adjusts the amplitude oscillations is attenuated. The correction takes the form:

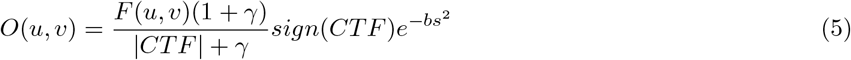

where *O*(*u, v*) and *F* (*u, v*) are respectively the Fourier transforms of the image and its underlying object, *s* is the magnitude of the coordinates *u* and *v* in frequency space, and *γ* is a positive regularization term that increases with spatial frequency so that phase flipping dominates at high resolution. Similar to Warp^22^, a low-pass filter is applied as part of the CTF correction with a default but adjustable B-factor, *b*, of 15 Å^−2^.

Extended Data Fig. 8 compares AreTomo3’s local CTF correction to the approach implemented in IsoNet^33^, which is based on Warp^22^. In contrast to AreTomo3, IsoNet applies a CTF deconvolution to the reconstructed tomograms rather than locally on the tilt-series and uses a single defocus value for the entire volume, which ignores the defocus gradient. A second key difference is the filter shape: IsoNet relies on a Wiener-like filter in contrast to the hybrid form described above. As a result, the drop in intensity at the zero-crossings in the Fourier domain is much more pronounced in the IsoNet case (Extended Data Fig. 8b-c). Though in principle there should be no information transfer at these spatial frequencies, in practice the intensity will be nonzero due to noise, and the presence of sharp features in Fourier space was anticipated to be disadvantageous for training denoising models.

### Comprehensive quality assessment

Every step of the cryoET workflow is susceptible to errors, which can propagate to and impair downstream steps. AreTomo3 provides a screening mechanism to exclude some low-quality data from subsequent processing. For each reconstruction, AreTomo3 records six metrics, spanning tomographic alignment, information transfer, and sample geometry. For alignment, it reports the tilt axis angle, maximum global shift, and the fraction of patches with failed local alignment. For information transfer, it provides the Thon ring resolution and CTF estimation score for the 0° image. Sample thickness is reported for geometry. For datasets collected from similar samples and with the same acquisition parameters, comparing these metrics offers insights into dataset quality (Extended Data Fig. 9). In addition to quality monitoring, these metrics can be leveraged for tomogram curation.

The effectiveness of applying *k*-means clustering to these metrics as a screening mechanism was evaluated by comparing its classification to visual inspection by expert annotators. A dataset of 789 tomograms of *Mycoplasma mycoides* JCVI-Syn3A “near-minimal” cells^34^ (minicells) was reviewed, with 245 high-quality tomograms retained by users for STA (Fig. 4a-b). *K*-means was then applied to independently classify the tomograms into two clusters. The cluster (Fig. 4c-h, cluster 0) with a higher standard deviation across all metrics was considered the rejected class. Compared to visual inspection, *k*-means clustering correctly classified 61% of user-rejected tomograms. Only 8% of user-selected tomograms were misclassified, even though a criterion during manual screening was ribosome abundance, which AreTomo3’s quality metrics do not directly report on.

**Figure 4:**
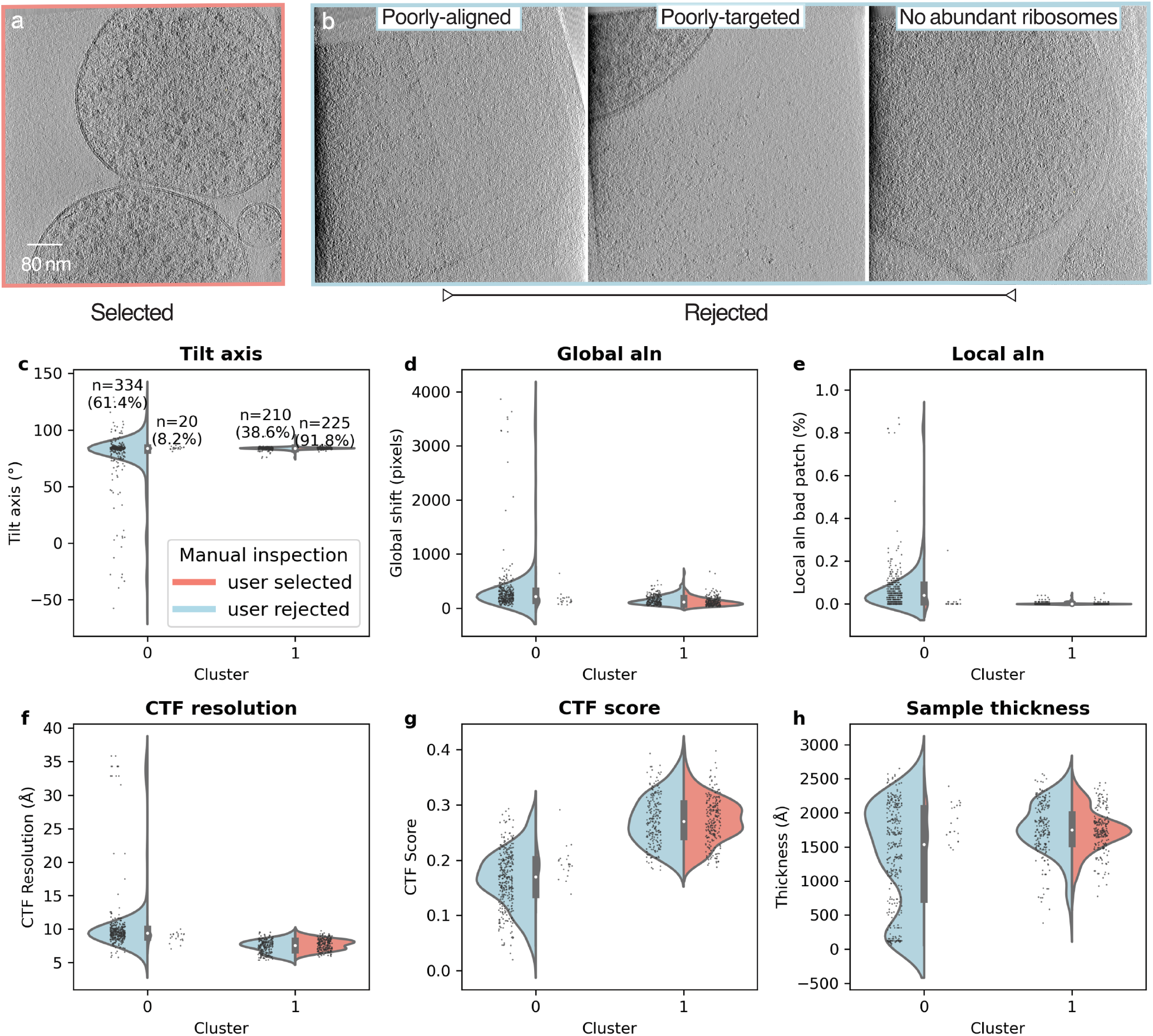
Visual inspection versus *k*-means clustering of AreTomo3’s metrics on minicell tomograms. During visual inspection, **a**. well-aligned tomograms with clearly discernible ribosomes were selected while **b**. tomograms that were poorly-aligned, mistargeted, and/or lacking abundant ribosomes were rejected. 245 of 789 tomograms were selected for downstream analysis based on these criteria. *K*-means clustering with 2 clusters was performed on AreTomo3’s metrics. The resulting clusters are color-coded by user annotation and plotted as violin plots with scattered data points overlaid for the following metrics: **c**. tilt axis angle, **d**. maximum global shift, **e**. percentage of patches with failed local alignment, **f**. Thon ring resolution, **g**. CTF estimation score for the zero-tilt image, and **h**. sample thickness. The number and percentage of user-rejected (blue) and user-selected (red) tomograms in each cluster are reported in panel **c**., with percentages calculated relative to the total user-rejected (544) and user-selected (245) tomograms. Compared to visual inspection, AreTomo3’s screening mechanism successfully classified over 61% of the user-rejected tomograms (334 out of 544) while misclassifying ∼8% of the user-selected tomograms (20 out of 245).

By comparison, a constant thresholding approach that classified tomograms based on specific cut-off values for all six of the reported metrics correctly classified 77% of the user-rejected tomograms but incorrectly rejected 26% of the user-selected tomogram, more than three times the rate of incorrect rejections of the *k*-means approach (Extended Data Fig. 10). The thresholds used for these metrics were largely chosen based on the dataset’s statistics in an effort to remove outliers. Though tuning the cut-off values might improve consistency with the results of visual inspection, such per-dataset adjustments are subjective and reduce the generalizability of this approach. To evaluate the generalizability of *k*-means clustering, this analysis was repeated for a phantom dataset^35^ and a dataset of affinity-captured lysosomes (Extended Data Fig. 11). In these cases, *k*-means clustering correctly classified 45–53% of the user-rejected cases into the rejected cluster for these datasets, with 13-25% of the user-selected tomograms misclassified. The reduced consistency between *k*-means clustering and user annotations for these datasets compared to the minicell dataset may reflect the more biologically-driven selection criteria that guided visual inspection, or the fact that the tomograms used for *k*-means clustering and visual inspection were reconstructed by different software packages. Regardless, the ability to automatically discard half of low-quality tomograms would significantly reduce the time spent on manual curation for large datasets.

### Contrast enhancement by 3D denoising

The second component of AreTomoLive, DenoisET, generates high-contrast tomograms, including during data collection. The basis for denoising is the ML algorithm Noise2Noise^18^, which learns to predict clean imaging data by training on paired noisy measurements of the same field-of-view but corrupted by independently and identically distributed noise. AreTomo3 generates the required training data by splitting the motion-corrected movies into even and odd frames, which are then summed and aligned using the same alignment parameters to reconstruct the corresponding even and odd tomograms. Although paired training data could be generated in other ways — for instance, by splitting the tilt-series into even and odd tilt images^36^ or the tomograms into every other slice along *z*^37^ — interleaved splitting at the frame level yields half-sets with the highest degree of structural similarity to best satisfy the premise of the Noise2Noise algorithm.

AreTomo3 further increases the spatial coherence of the training data through its CTF correction. In addition to correcting the CTF, this deconvolution applies a low-pass filter to focus the representative capacity of the neural network on the low to intermediate resolution features responsible for contrast. The impact of this CTF correction on subsequent denoising was compared to the approach taken by IsoNet^33^ and Warp^22^, which deconvolves the CTF from the tomograms rather than the tilt-series using one defocus value for the entire volume. We found that a stronger low-pass filter was needed to achieve the highest quality denoising in the IsoNet/Warp case compared to AreTomo3’s default filter strength (Extended Data Fig. 8). However, applying a strong low-pass filter during data preprocessing blurs intermediate resolution features that ideally the neural network would learn during training and retain during inference. Consistent with this, we observed that intermediate resolution features were better preserved after denoising tomograms CTF-corrected by AreTomo3 compared to IsoNet across different specimens, including a lamella of a cilium and centrosome and a phantom dataset^35^ (Fig. 5 and Extended Data Fig. 12).

**Figure 5:**
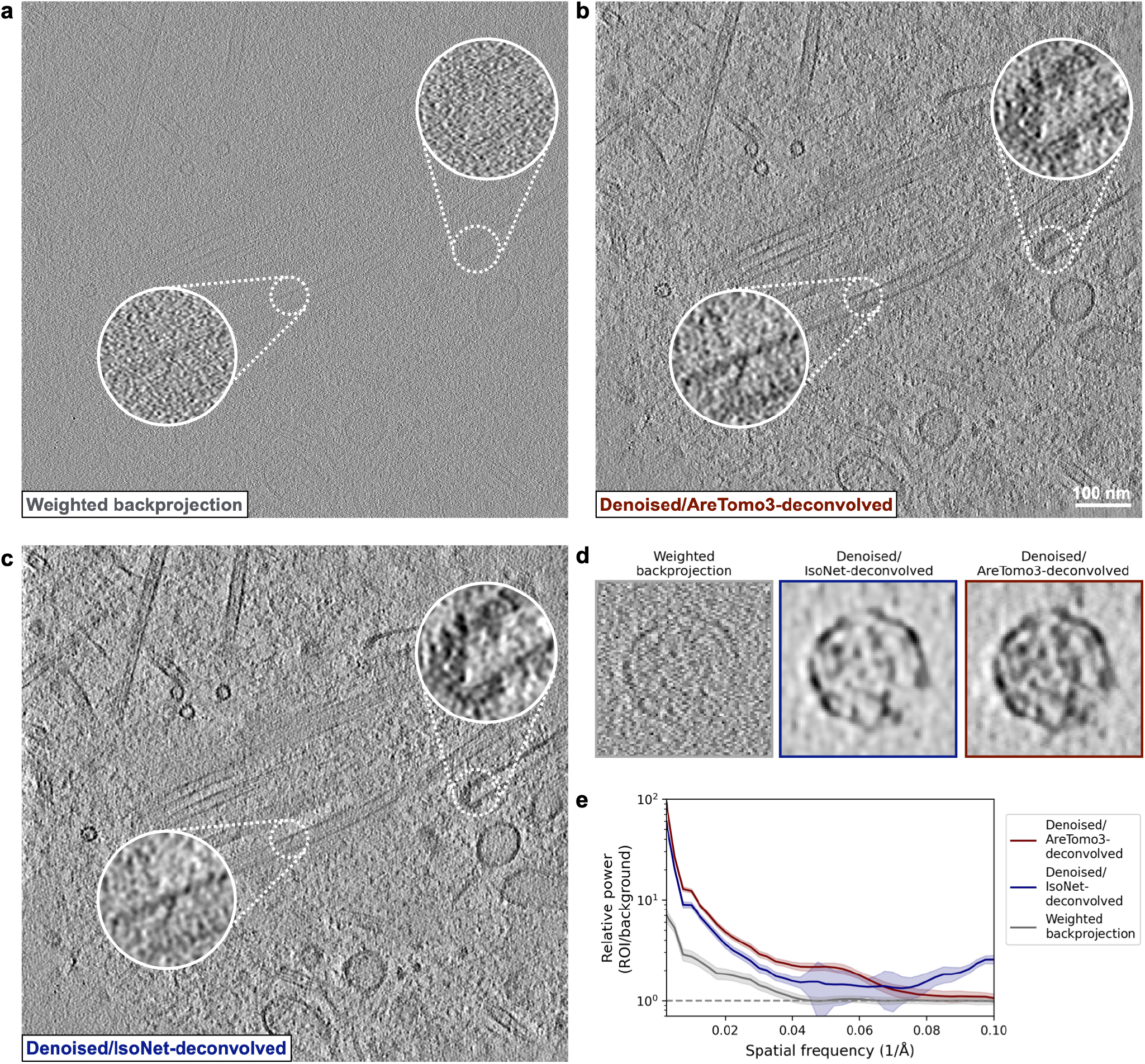
AreTomo3’s CTF correction improves the retention of intermediate resolution features during denoising. A 10 Å slice from a lamella of a cilium and centrosome is compared from the **a**. undenoised tomogram and after denoising the **b**. AreTomo3-corrected versus **c**. IsoNet-corrected tomograms. The insets highlight proteins embedded in the membrane and arrayed along the filament. **d**. Representative virus-like-particles from a phantom dataset^35^ are compared from the indicated tomograms. **e**. Contrast enhancement was estimated by computing the ratio of the power spectra of these regions of interest (ROI) to background regions for the indicated tomogram type, with the standard deviation indicated by shading.

Contrast enhancement was quantified by computing the ratio of the power spectra of regions of interest to background regions in the phantom dataset. From this analysis, we estimated that denoising the AreTomo3-corrected tomograms on average provided a 5-fold increase in contrast out to the first zero-crossing relative to the weighted backprojection volumes, and a 1.5-fold increase compared to denoising the IsoNet-corrected tomograms. Critically, contrast enhancement extended to intermediate resolution without introducing artifacts at high resolution as observed in the IsoNet case (Fig. 5d-e).

In addition to its CTF correction, DenoisET leverages AreTomo3’s quality metrics to improve the effectiveness of denoising (Extended Data Fig. 13). These metrics are used to automate the selection of training data by retaining only the tomograms that meet specific thresholds for the six reported metrics, yielding more stringent curation than *k*-means clustering. Across diverse datasets, we have found that 20-40 high-quality tomograms generally suffice for robust model training. For denoising in real-time, DenoisET initiates training once sufficient high-quality tomograms are detected. Training is automatically terminated based on the emergence of checkerboard artifacts in the denoised tomograms, which is more sensitive to overdenoising than other metrics like the training loss (Extended Data Fig. 14). Automating this step is critical since visual inspection to select the optimal training epoch^37,38^ is unviable for real-time processing, and the number of training epochs that yields the best denoising is unpredictable due to the stochastic nature of model training and the high variability of cryoET data. Once training finishes, DenoisET performs inference on all available tomograms, including those excluded from the training set, and transitions to monitoring for additional CTF-corrected tomograms to denoise. DenoisET can also be run in an inference-only mode when a suitable pre-trained model is available and provides several pre-trained models used to provide denoised tomograms for the CryoET Data Portal^39^. However, we have often found it advantageous to train a new model from scratch due to the limited generalizability of pretrained models (Extended Data Fig. 15) and given that the time and computational cost required to train a new denoising model are not prohibitive.

## Discussion

Here we present AreTomoLive, an accelerated preprocessing pipeline that automates tomographic alignment, reconstruction, and contrast enhancement. These tasks are accomplished by its constituent programs, AreTomo3 and DenoisET, which are designed to run in parallel to deliver real-time feedback on data quality and the processed data and metadata required for downstream analysis. Beyond seamlessly integrating the widely-used MotionCor2^16^ and AreTomo^17^ packages, AreTomo3 introduces several new features that collectively enhance alignment and reconstruction accuracy. The resulting increase in local resolution in the tomograms is expected to benefit particle picking and segmentation in addition to traditional STA algorithms that rely on 3D subtomograms^40,41^. However, the RELION-4/5 STA workflows^21,42^ that extract data from tilts-series in a pseudo-subtomogram approach currently exploit only AreTomo3’s global and not its local BIM corrections, nor its correction of the *α*_0_ offset. Future efforts will explore how to bridge this gap, either by defining a common deformation model that maps coordinates from locally-aligned and *α*_0_-corrected tomograms to the tilt-series or by directing AreTomo3 to generate the per-particle tilt-series used as pseudo-subtomograms during STA. This approach would leverage AreTomo3’s rigorous local alignments as a more effective starting point for subtomogram polishing.

Despite these improvements to alignment and reconstruction, there will invariably be low-quality tomograms due to mistargeting, ice contamination, and other issues that cannot be algorithmically resolved. Excluding these tomograms from downstream processing is critical to ensure high quality data both to train ML algorithms and for high-resolution STA. To address the bottleneck of data curation, AreTomo3 tracks quality metrics that report on sample geometry, alignment quality, and information transfer and provides auxiliary tools to filter out low-quality tomograms using *k*-means clustering. Compared to applying constant thresholds, this approach partitions tomograms more similarly to expert annotations and requires minimal parameter tuning from one dataset to the next. In contrast to supervised deep learning methods, this unsupervised approach avoids the need for large manually-annotated datasets. Including additional metrics — in particular, the *α*_0_ and *β*_0_ offsets for lamella samples — and different clustering methods may further improve the effectiveness of this screening strategy.

Like data curation, contrast enhancement is often a critical post-reconstruction task. Though Noise2Noise has previously been applied to cryoET data^36,43^, leveraging AreTomo3’s local CTF correction and quality metrics increases the quality of denoising. This improvement is particularly evident in the increased preservation of intermediate resolution features, which should enable better discrimination between densities of similar size during particle picking. DenoisET integrates this denoising scheme into AreTomoLive to automate contrast enhancement by algorithmically selecting training data and transitioning from training to inference without the need for visual inspection. Based on advances in the ML field since Noise2Noise was first described, it will be useful to explore whether compound loss functions or different neural network architectures further improve denoising^44,45^. However, this workflow already achieves significant contrast enhancement that could be used to refine tomographic alignment. Specifically, we envision forward-projecting the denoised tomograms and using the resulting contrast-enhanced tilt-series to refine the local alignments and tilt angles under a Bayesian framework. The increased alignment accuracy from this subvolume polishing scheme would yield more spatially-coherent tomograms, which in turn would provide the basis for superior denoising.

Increasing throughput is critical to realize the full potential of cryoET, from STA of challenging molecular targets to training generalizable ML models and robust statistical analysis of cellular ultrastructure. AreTomoLive responds to this need by delivering an automated pipeline that keeps pace with data collection to provide users with immediate feedback on data quality and the tomograms, tilt-series, and metadata required for downstream processing tasks. This pipeline is currently being used to reprocess ∼17,000 tilt-series on the recently released CryoET Data Portal^39^ to supply the community with well-aligned tomograms in a standardized format, demonstrating this pipeline’s effectiveness across diverse datasets and capacity for throughput. AreTomoLive is also designed with the flexibility to harness additional GPU resources to ensure that preprocessing can continue to scale with data collection as more computationally-intensive algorithms are developed. Continued efforts to improve the efficiency and robustness of every aspect of data processing will play a critical role in transforming cryoET from a niche technique into a routine method for *in situ* structural biology.

## Methods

### Sample preparation

For the minicell samples, JCVI-syn3A cells (J. Craig Venter Institute) were grown at 37°C without aeration or agitation in SP4-KO medium until reaching an A600 of 0.11. 4 *µ*L of cell supernatant was then applied directly to 200-mesh Cu R2/1 Quantifoil grids (Quantifoil). For the purified synaptosome samples, hippocampus tissue from 10 week old rats (Transnetyx) was homogenized by mortar and pestle, followed by 10 minutes of centrifugation at 800 rpm at 4°C. Samples were diluted in Hibernate A media (Transnetyx) at a ratio of 1:10 or 1:50 before being applied to 200-mesh Cu R1.2/1.3 or R2/2 Quantifoil grids (Quantifoil).

For the affinity-captured lysosomes, a HEK293T cell line with a knock-in mEGFP tag on the C-terminus of TMEM192 was used. Cells were lysed in a hypotonic homogenization buffer (25 mM Tris-HCL pH 7.5, 50 mM sucrose, 0.2 mM EGTA, 0.5 mM MgCl_2_) and sheared with a 23G syringe. The lysate was equilibrated in sucrose buffer (2.5 M sucrose, 0.2 mM EGTA, 0.5 mM MgCl_2_), and the nuclear fraction was removed by centrifugation at 1000×g for 10 minutes. For the gold-labelled samples, lysate was incubated with 0.5-2 *µ*M SidK-HaloTag or SpyCatcher-HaloTag fusion protein conjugated to 2.2 or 3 nm maleimide-functionalized gold particles (Nanopartz), respectively. Incubation was performed in a ThermoMixer (Eppendorf) at 4°C for 1 hour with shaking at 500 rpm. 6 *µ*L of lysate was then applied to grids functionalized with anti-GFP nanobodies^46^, incubated for 10 seconds, and washed with 6 *µ*L of phosphate buffer saline (PBS). This sequence was repeated three times, with excess liquid removed by filter paper after each addition and incubation. The grid was then washed with 6 *µ*L of PBS, and the addition of lysate and PBS wash steps were repeated once more. A final 6 *µ*L of PBS were added to the grid.

The minicell, synaptosome, and lysosome samples described above were blotted using a Whitman #1 blotting paper for 4-6 seconds with the chamber conditions set to 4°C and 95-100% humidity and plunge-frozen in liquid ethane at -180°C using a Leica GP2.

For the cilia and centrosome lamellae, hTERT RPE-1 cells (ATCC CRL-4000, female) stably expressing HTR6-HaloTag3^47^ were plated at ∼20,000 cells/cm^2^ on glow-discharged R2/2 silicone dioxide grids with gold bars (Quantifoil) in 35 mm glass-bottom dish (20 mm #1.5 coverglass, Cellvis) in 10% serum-containing media (DMEM:F12, ATCC 30-2006; Day 0) at 37°C in 5% CO2. The next day, cells were serum-deprived with 0% FBS media and 100 ng/ml doxycycline to induce HTR6-HaloTag expression. After 48 hours of serum deprivation, HTR6-HaloTag cells were labeled with 250 nM Janelia Fluor 552 (JF552) dye for 1 hour. Cells were rinsed in PBS without added calcium and placed in a HEPES-buffered imaging media (140 mM NaCl, 20 mM HEPES, 2.5 mM KCl, 1.8 mM CaCl_2_, 1.0 mM MgCl_2_, pH 7.4; Live Cell Imaging Solution, TFS). After a 5 second incubation in imaging media with 10% glycerol, the grids were plunge-frozen on a Leica EM GP2 plunge freezer with the chamber temperature set to 35°C and humidity at 95% and blotted for 3 seconds. Lamella were generated using fluorescence-guided milling on a Thermo Aquilos with an integrated fluorescence microscope (iFLM).

For the yeast lamellae, a *S. cerevisiae* strain expressing Myo5-GFP, Myo3-GFP, and Ab1-RFP fusion proteins were grown to log phase, applied to 200-mesh Cu R3.5/1 grids, and plunge frozen on a Leica GP24. Lamella milling was performed in an Arctic FIB-SEM (TFS) under cryogenic conditions, and the TFS WebUI software was used for the entire workflow. Grids were coated with three platinum layers: a sputtering layer (12 kV, 120 nA, 120 seconds) of metallic platinum, a gas injection system layer (120 seconds) of organo-platinum, and a second sputtering layer (12 kV, 120 nA, 120 seconds) of metallic platinum. Lamellae were milled in 4 sequential steps using a 30 kV Xenon ion beam and 1000 pA, 300 pA, 100 pA, and 30 pA respectively for the rough, medium, fine, and polishing steps.

### Data collection

All data were collected on a Krios G4 (TFS) equipped with an X-FEG electron gun, a Falcon 4i detector, and the SelectrisX energy filter. The pixel size was set to 1.54 Å for the minicell, synaptosome, and lysosome samples; 2.37 Å for the cilia and centrosome lamellae; and 1.94 Å for the yeast lamellae. A total dose of 120 ^−^/Å^2^ was applied for the lysosome, synaptosome, and lamella data, and 150 e^−^/Å^2^ was applied for the minicell data. The target defocus was 2 *µ*M for the synaptosome, lysosome, minicell, cilia and centrosome lamellae; and 2.5-4 *µ*M for the yeast lamellae. The acquisition software was the Tomography 5 software (TFS), with beam-image shifts used to collect multiple targets at each stage position. Movies were saved in EER format.

### Self-adaptive defocus search range and step size during tilt-series CTF estimation

AreTomoLive uses Eq. 1 to automatically decide the defocus search range and step in its CTF estimation at different magnifications. The formula for the CTF phase shift (*ϕ*) is given by:

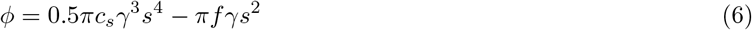

where *γ* and *s* are the electron wavelength and the spatial frequency, respectively. As can be seen, the spherical aberration contributes much less to the phase shift than the defocus due to the product of *γ*^3^ and *s*^4^. When the contribution of the spherical aberration is ignored, two CTFs have approximately the same phase shifts if Eq. 7 is satisfied:

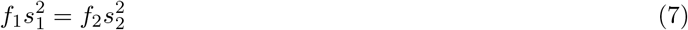

The discrete spatial frequency *s*^(*i*)^ can be expressed in Eq. 8 where *p* denotes the pixel size, and *n* and *N* refer to the *n*^*th*^ Fourier component and the size of the Fourier transform, respectively.

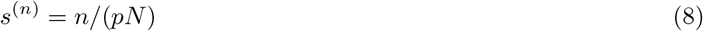

Plugging Eq. 8 into Eq. 7 gives:

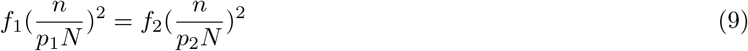

Eq. 1 can thus be obtained by simplifying Eq. 9. Another application of Eq. 1 is to help users choose an optimal defocus setting for data collection at different magnifications. For example, the CTF for 4 *µ*m defocus at 2 Å pixel size has approximately the same phase shift as that for 1 *µ*m defocus at 1 Å pixel size.

### Defocus distribution within tilt images

Extended Data Fig. 2 shows a tomogram in which sample tilt was observed in both the *xz* and *yz* planes, indicating both nonzero *α*_0_ and *β*_0_. This observation motivated our derivation of Eq. 2 to model the defocus distribution within tilt images by taking into account these two angular offsets.

Ignoring sample thickness, Extended Data Fig. 2 presents two diagrams that show a pre-tilted sample at 0° and-*α* stage positions, respectively. For a given point at (Δ*x*, Δ*y*) in the projection plane, Extended Data Fig. 2a shows that Δ*z*, the difference in the *z* coordinate of the corresponding sample point labeled with a dark dot between the two positions, is nonzero. Δ*z* can be formulated based on the tilting geometry as:

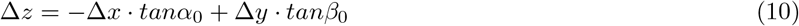

Eq. 10 shows that *α*_0_ and *β*_0_ both contribute to the *z* height change at any point. When the sample is tilted by -*α* as shown in Extended Data Fig. 2b, the new tilting geometry can be formulated as:

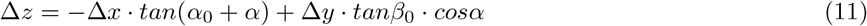

While the Δ*x* term in Eq. 11 is straightforward, it is worth noting that tilt reduces the effect of *β*_0_ on inducing the sample *z*-height change. If a sample were able to be tilted to 90°, *β*_0_ would not introduce any change in *z*-height. The most noticeable effect of *β*_0_ is therefore at low tilt angles. This information is incorporated into our optimization process. Note that (Δ*x*, Δ*y*) is relative to the aligned coordinate system, whereas (Δ*x*’, Δ*y*’) in Eq. 2 is with respect to the raw image coordinate system. Eq. 2 can be derived from Eq. 10 by applying the rotation matrix associated with the tilt axis angle.

### Automation of 3D denoising

When run in parallel with AreTomo3 during live data collection, DenoisET continually monitors the quality metrics file generated by AreTomo3 to track tomograms with quality scores that meet all of the specified thresholds. The metrics currently considered are sample thickness, the tilt axis angle, the maximum global shift of the untilted image, the fraction of bad local patches, and the CTF resolution and score at 0°. The default thresholds for each of these metrics can be changed by users. Training begins as soon as sufficient high-quality tomograms are detected. If the number of high-quality tomograms exceeds a threshold value, the selected tomograms will be sorted by their global shift parameter and the top-scoring tomograms will be used for training. For offline processing, an AreTomo3 metrics file, a list of training tomograms, or a folder containing tomograms can be provided. In general, we have found that 20-40 tomograms suffice for model training. Paired subvolumes of 96 pixels cubed are extracted from the corresponding even and odd tomograms, with data augmentation (consisting of random 90° rotations, mirror operations across the center plane, and swapping which subvolume is considered even or odd) performed.

DenoisET implements the Noise2Noise algorithm^18^ for cryoET data, with a similar model architecture as implemented in Topaz-Denoise^43^ but a kernel size of 3 pixels for all layers. In addition to the training and validation loss scores, DenoisET tracks the standard deviation of denoised subvolumes and a metric that quantifies the emergence of checkerboard artifacts during each training epoch. Due to GPU memory constraints, the denoising model is applied to overlapping subvolumes with the padded regions excised during stitching to avoid boundary artifacts. However, empirically we have observed that checkerboard features tend to emerge concurrently with other overdenoising artifacts, such as halos around the perimeter of high contrast regions and oversuppression of low contrast features. To quantify this checkerboard artifact, the intensity difference between adjacent pixels in the *xy* planes that form the border between neighboring subvolumes is compared to this value from nearby adjacent planes that lie entirely within each subvolume under consideration. The relative change is computed for each pair of subvolumes with neighboring *xy* planes within the full denoised volume. Border regions for the other planes are not considered to avoid missing wedge artifacts. When the mean of this statistic exceeds a threshold value, DenoisET automatically reloads the model from the prior epoch and transitions from training to inference. If the checkerboard metric does not exceed this threshold value during training, DenoisET selects the model from the training epoch associated with the highest standard deviation for the denoised subvolumes and uses that to perform inference. For real-time processing, DenoisET continually checks for new tomograms to denoise during the inference phase.

### Evaluation of contrast enhancement

The impact of AreTomo3’s CTF correction on denoising was assessed by comparison with the deconvolution implemented in IsoNet^33^, which is based on the approach taken in Warp^22^. The IsoNet CTF deconvolution was applied to weighted backprojection tomograms reconstructed by AreTomo3 using the defocus value estimated at 0°. A grid search was performed to determine optimal values for the fall-off and strength parameters in the SSNR term of the Wiener-like filter. A denoising model was trained separately on each set of tomograms, and the combination of values that led to the most effective denoising based on visual inspection was chosen. Empirically we found that denoising was more sensitive to the fall-off than the strength parameter. No tuning of the B-factor that adjusts the strength of AreTomo3’s CTF deconvolution was performed for any of the datasets presented in this paper. A separate denoising model was trained on the AreTomo3-deconvolved tomograms.

Contrast enhancement was quantified by computing the ratio of the power spectra of regions of interest to background regions for different tomogram types. The regions of interest consisted of 224 virus-like-particles identified across 45 tomograms, while 135 background regions were selected from the same set of tomograms based on local intensity statistics. 80^3^ pixel subvolumes centered on each region of interest and background region were extracted from tomograms reconstructed with a pixel size of 5 Å. A cosine mask was applied to each subvolume in real space prior to computing the radially-averaged power spectrum. The ratio of the mean power spectra of the regions of interest and background regions was then calculated. Comparing the power of regions of interest and background from the same tomogram type provides an estimate of the inherent contrast of localized regions that would be targeted by downstream analysis such as particle picking. Importantly, this ratio will only increase if contrast enhancement is more pronounced in the regions of interest; strategies that uniformly increase contrast throughout the tomogram will not affect this ratio.

## Data Availability

The minicell, synaptosome, and phantom tomograms are available on the CryoET Data Portal under deposition IDs CZCDP-10312, CZCDP-10313, and CZCDP-10310, respectively.

## Code Availability

AreTomo3 is open-source under the BSD-3-Clause license, and its source code can be downloaded from //github.com/czimaginginstitute/AreTomo3. DenoisET is open-source under the MIT license and available at https://github.com/apeck12/denoiset.

## Acknowledgments

Throughout the history of AreTomo3’s development, the authors have constantly received encouragement, feedback, and suggestions from the cryoET community. Their enthusiastic support is greatly appreciated. We also acknowledge the University of California San Francisco and Howard Hughes Medical Institute for agreeing to make MotionCor2 and AreTomo open-source, which expedited AreTomo3 development. We thank Montserrat Barcena (Leiden University Medical Center) for sharing the tilt-series of arterivirus-infected cells to assist code development. We also thank Momoko Shiozaki (Janelia Research Campus) for assistance with lamella preparation for the cilia sample and Hang Cheng for providing the yeast samples. The CZ Imaging Institute is made possible with support from the Chan Zuckerberg Initiative (CZII-2023–327779).

## Author contributions

A.P. designed and implemented DenoisET for 3D denoising of tomograms; M.P. and S.Z. contributed to its development. S.Z. designed and implemented AreTomo3; A.P. contributed to the development. A.P., D.K., B.C., and S.Z. carried out the estimation of contrast enhancement. Y.Y., A.P., C.P., and S.Z. designed and implemented the module for automated quality assessment. Y.Y., D.S., and M.P. performed visual inspection of the tomograms for quality assessment. M.P., Y.Y, D.S., and B.S. estimated throughput enhancement. M.P. and Y.Y. conducted thorough tests of the pipeline. U.E., J.H., Y.Y., and S.Z analyzed the reconstruction handedness. C.P. designed the party hat plot for real-time performance analysis and S.Z. contributed to the design. M.P, H.S., N.S.H., H.S., D.S., J.P., G.G., and S.S. prepared the samples, and M.P., D.S., and L.M. collected the datasets analyzed in this work. S.Z. and D.A. designed the architecture of AreTomoLive. B.C., C.P., and D.A. provided advice and guidance throughout the entire development process. B.C., C.P., D.A., and S.Z. conceived this project. Everyone contributed to writing this manuscript.

