## Supplementary Figures for "AreTomoLive: Automated reconstruction of comprehensively-corrected and denoised cryo-electron tomograms in real-time and at high throughput"

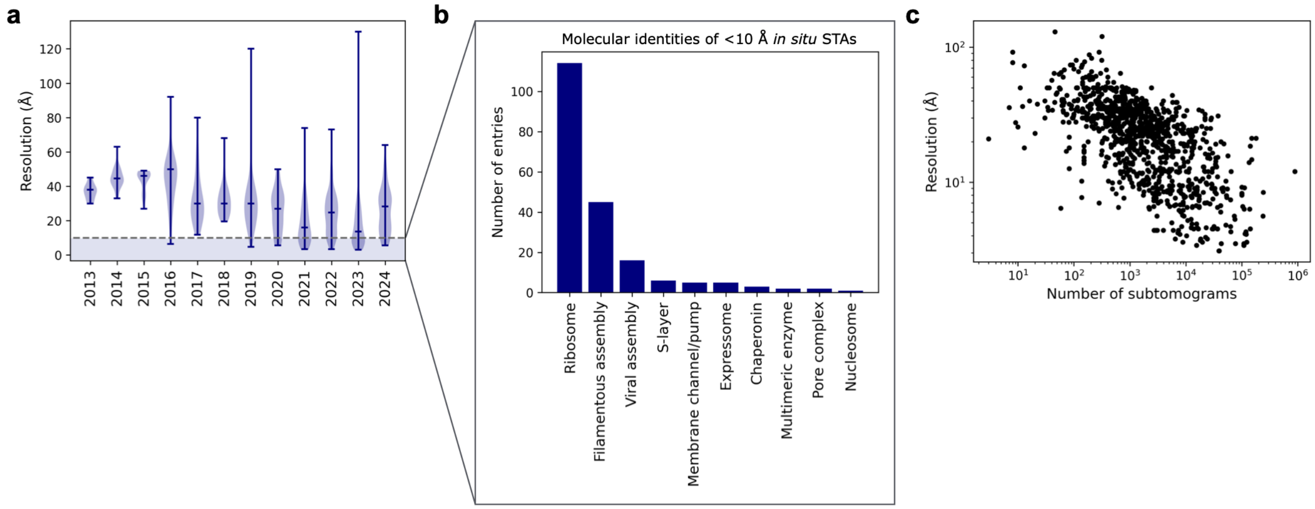


**Extended Data Figure** 1: **Characteristics of *in situ* subtomogram averages deposited in the Electron Microscopy Data Bank.** Structures deposited prior to October 2024 were included in the analysis if they were determined by subtomogram averaging from cell or tissue samples. **a.** The resolution of these structures is shown as a function of year, with the dashed line indicating 10 Å. **b.** The molecular identities of subnanometer resolution structures are categorized. Ribosomes, which are highly abundant, and viral and filamentous assemblies, which often have high symmetry, collectively account for nearly 90% of these sub-10 Å structures. **c.** Each structure's reported resolution is plotted as a function of the number of subtomograms used for averaging.

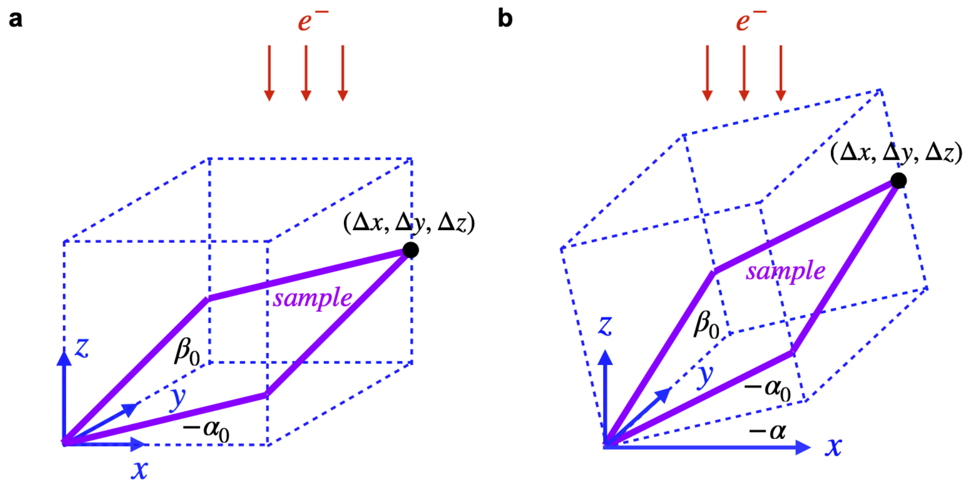


**Extended Data Figure** 2: **Tilting geometry of a sample with nonzero $\alpha_0$ and $\beta_0$ tilt offsets.** The sample is represented by the purple parallelograms. The boxes outlined in dashed lines indicate the initial sample tilting geometry prior to accounting for the tilt offsets. The $y$-axis is the tilt axis. **a.** The sample is at the stage position corresponding to 0° tilt. **b.** The stage is tilted by -$\alpha$ degrees.

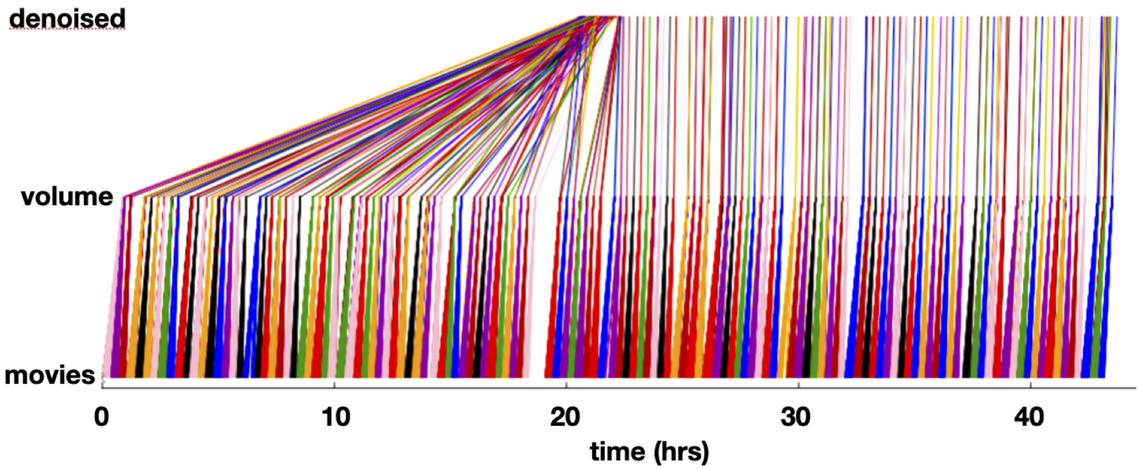


**Extended Data Figure** 3: **A "party hat plot" showing that AreTomoLive achieved real time tomographic reconstruction and 3D denoising.** This plot has three axes that show the relative timestamps of when the raw tilt-series movies, aligned and corrected tomograms, and denoised volumes were saved to disk across a multi-day data collection session. Since the colored lines are nearly vertical, it means that tomographic reconstruction and 3D denoising ran nearly as fast as data collection. The entire dataset contains 267 tilt series collected with a tilt-range of $\pm 45^{\circ}$ and a $3^{\circ}$ angular step.

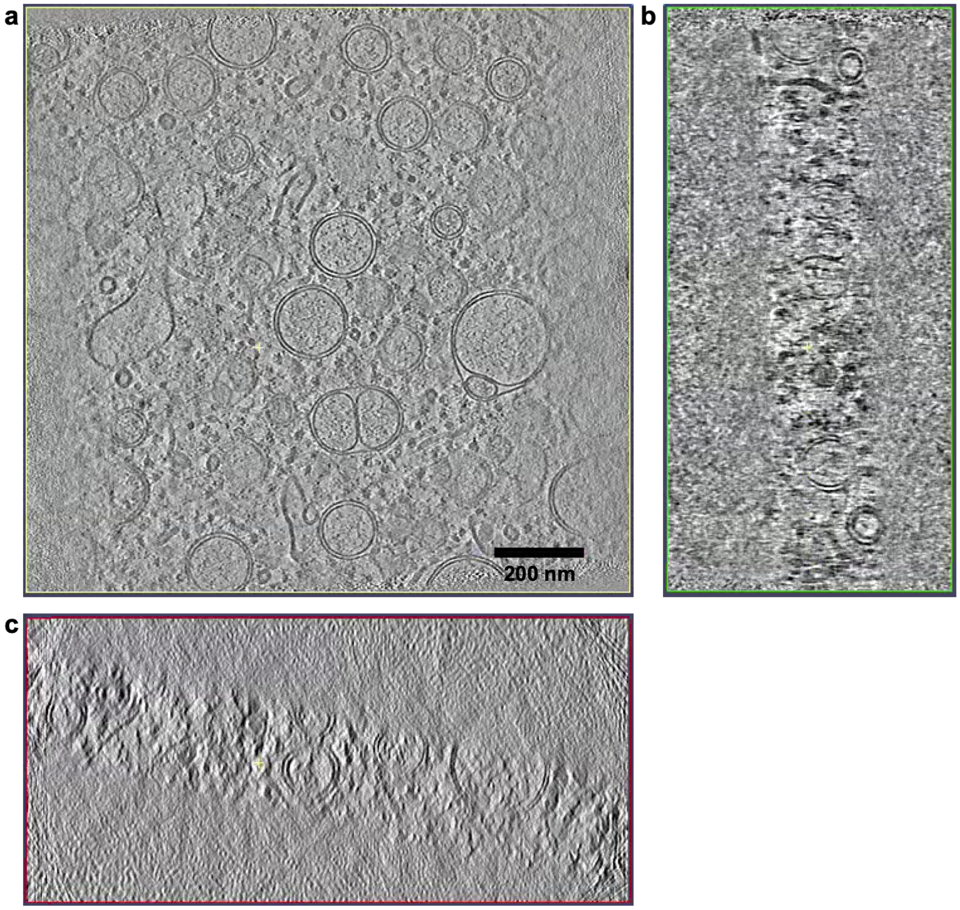


**Extended Data Figure** 4: **Three orthogonal tomographic slices from a tomogram of a FIB-milled lamella of an arterivirus-infected cell. a.** A slice from the $xy$ plane. **b.** A slice from the $yz$ plane shows a noticeable $\beta_0$ tilt. The estimate by AreTomo3 is -2.2°. **c.** A slice from the $xz$ plane shows a significant $\alpha_0$ tilt. The estimate by AreTomo3 is 12.3°.

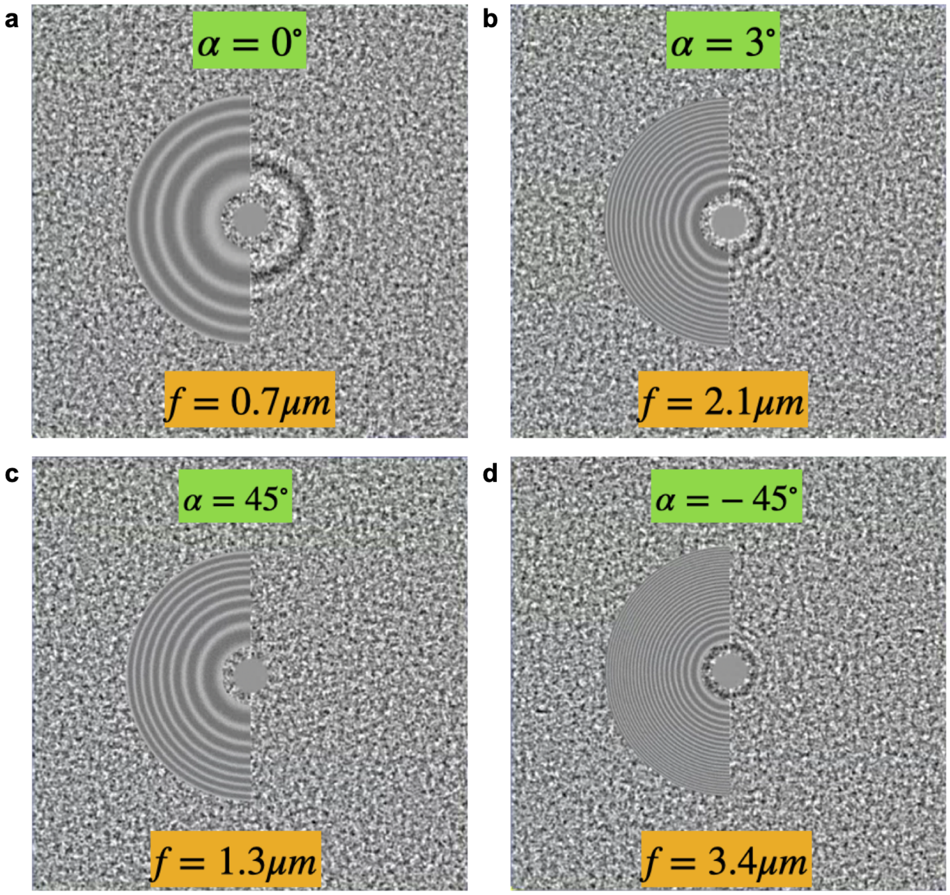


**Extended Data Figure** 5: **CTF estimation on a tilt-series of affinity-captured lysosomes collected according to a modified TYGRESS[25] scheme.** This tilt-series was collected with a 1.54 Å pixel size, and 10-fold more electron dose was applied to the 0° image than the tilted images. AreTomoLive was able to reliably estimate the CTF parameters despite the dose discontinuity and high defocus fluctuation from tilt to tilt. **a.** The tilt image at 0° was collected at a much lower defocus setting to facilitate 2D template matching. **b.** The tilt image at 3° was collected at a much higher defocus setting than that at 0° to increase contrast. **c.** The tilt image collected at 45° shows almost no visible Thon rings. Despite this, AreTomoLive delivered a reasonable estimate. **d.** The tilt image collected at -45° was found to have much higher defocus than that at 45°, an asymmetry that indicates an autofocusing error during data collection.

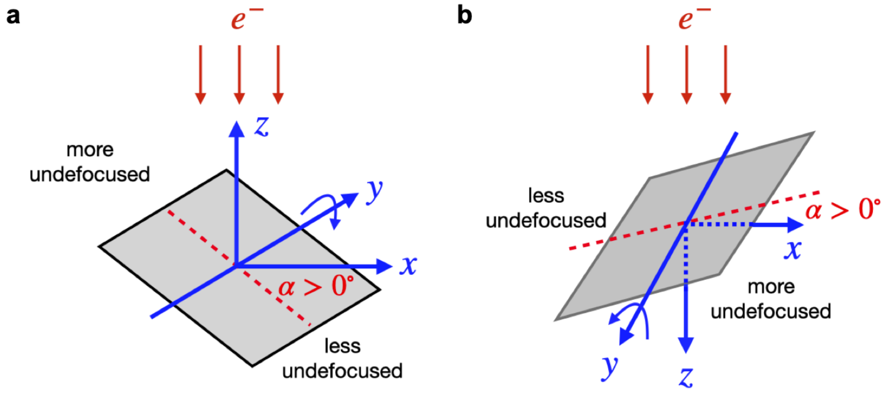


**Extended Data Figure** 6: **Two commonly used coordinate systems and their defocus handedness. a.** The coordinate system associated with positive defocus handedness, with the $z$-axis pointing to the electron source. **b.** The coordinate system associated with negative defocus handedness, with the $z$-axis pointing away from the electron source. In both coordinate systems, the $y$-axis corresponds to the sample rotation axis.

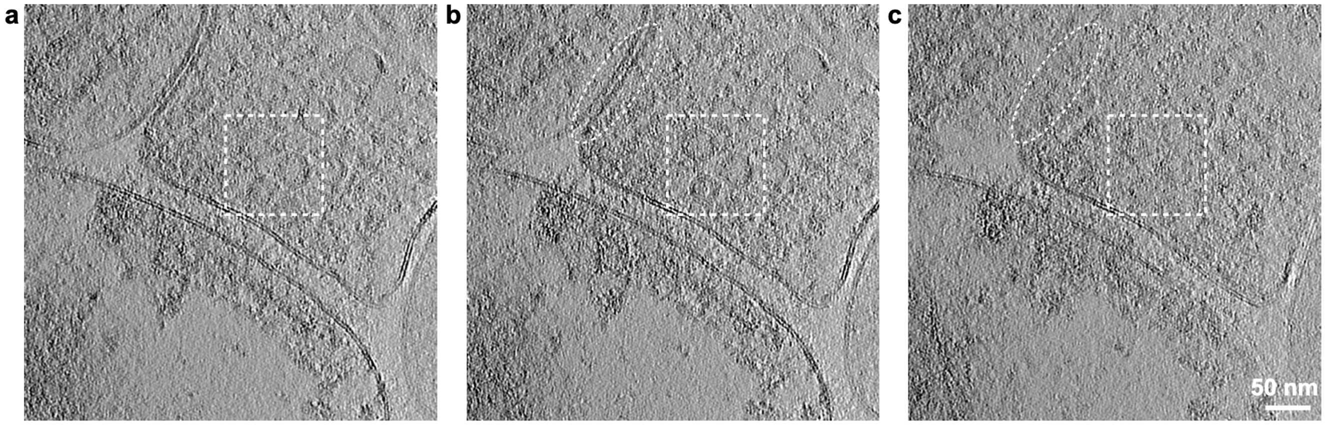


**Extended Data Figure 7**: **Correct estimation of sample thickness improves tomographic alignment.** The same tilt-series was aligned with three different values for the sample thickness used during the alignment process: **a.** 770 Å; **b.** 1500 Å, which is the sample thickness estimated by AreTomoLive; and **c.** 3800 Å. The presynaptic vesicles outlined in the square box are most distinct in **b.** The membranes labeled inside the oval in **b** are also better defined than in **c.**

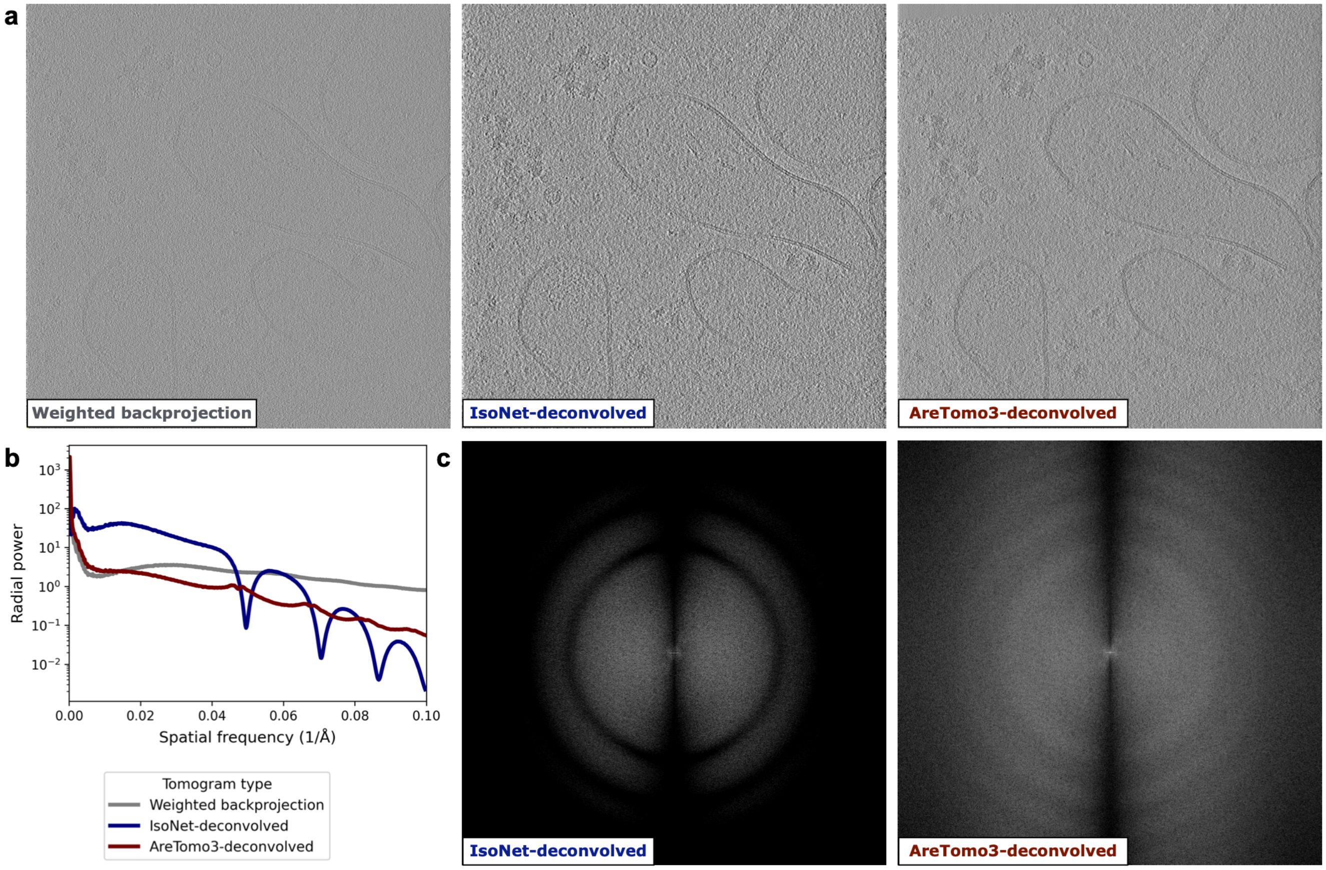


**Extended Data Figure** 8: **Comparison of AreTomo3's and IsoNet's[33] CTF deconvolution. a.** A tomogram reconstructed by weighted backprojection (left) from a phantom dataset[35] was CTF-deconvolved by IsoNet (center) or AreTomo3 (right). IsoNet applies a Wiener filter-like deconvolution post-reconstruction using a single defocus value for the full volume, whereas AreTomo3 applies a hybrid deconvolution locally to the tilt-series during the reconstruction process to account for the defocus gradient. **b.** The radial intensity profiles of the Fourier transform of each tomogram are compared. **c.** The Fourier transform through the CTF-deconvolved tomograms visualized in **a.** are compared.

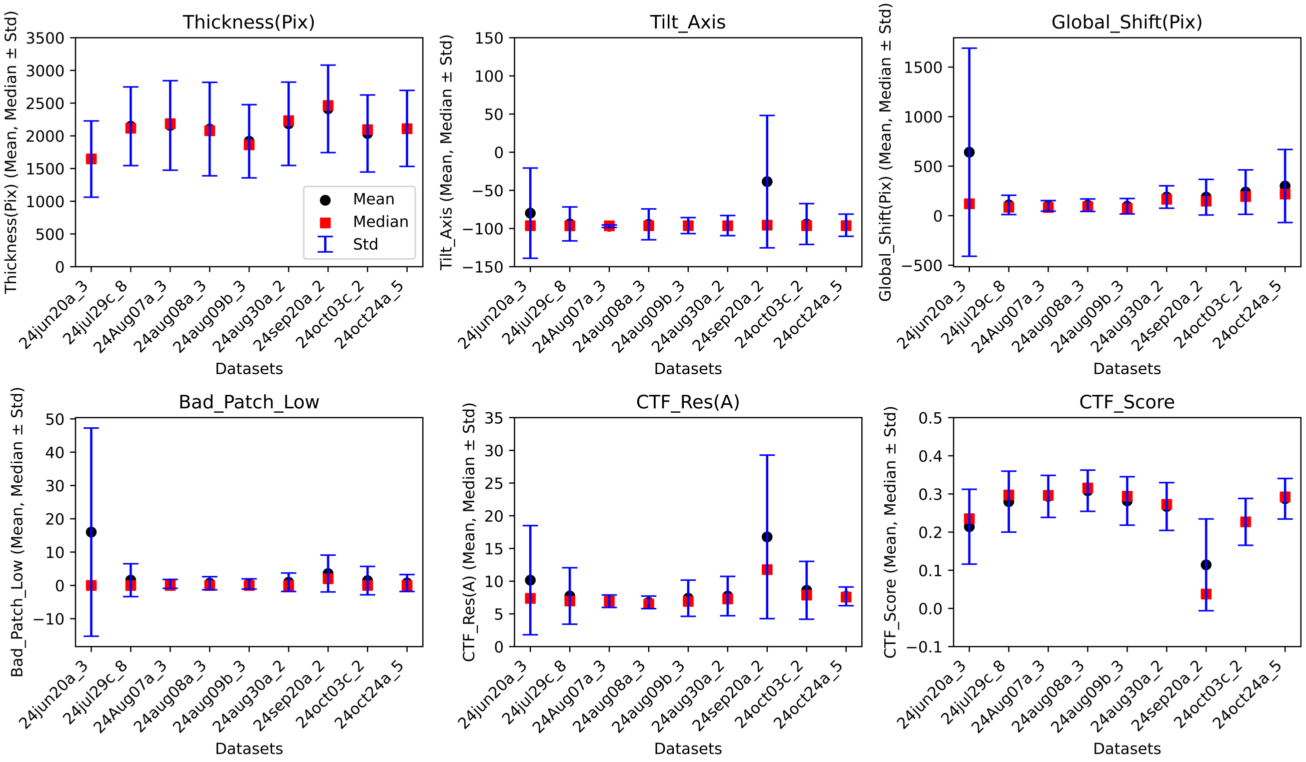


**Extended Data Figure 9: Comparison of AreTomo3's metrics across data collection sessions with the same acquisition parameters and of similar samples for quality monitoring.** AreTomo3's metrics for eight datasets acquired from affinity-captured lysosome samples using identical optical settings and acquisition schemes are shown. Outliers, particularly in the estimated tilt axis angle and CTF scores, often indicate low-quality data. Providing this information in real time during data acquisition can help identify the need for adjustments to data collection or processing parameters.

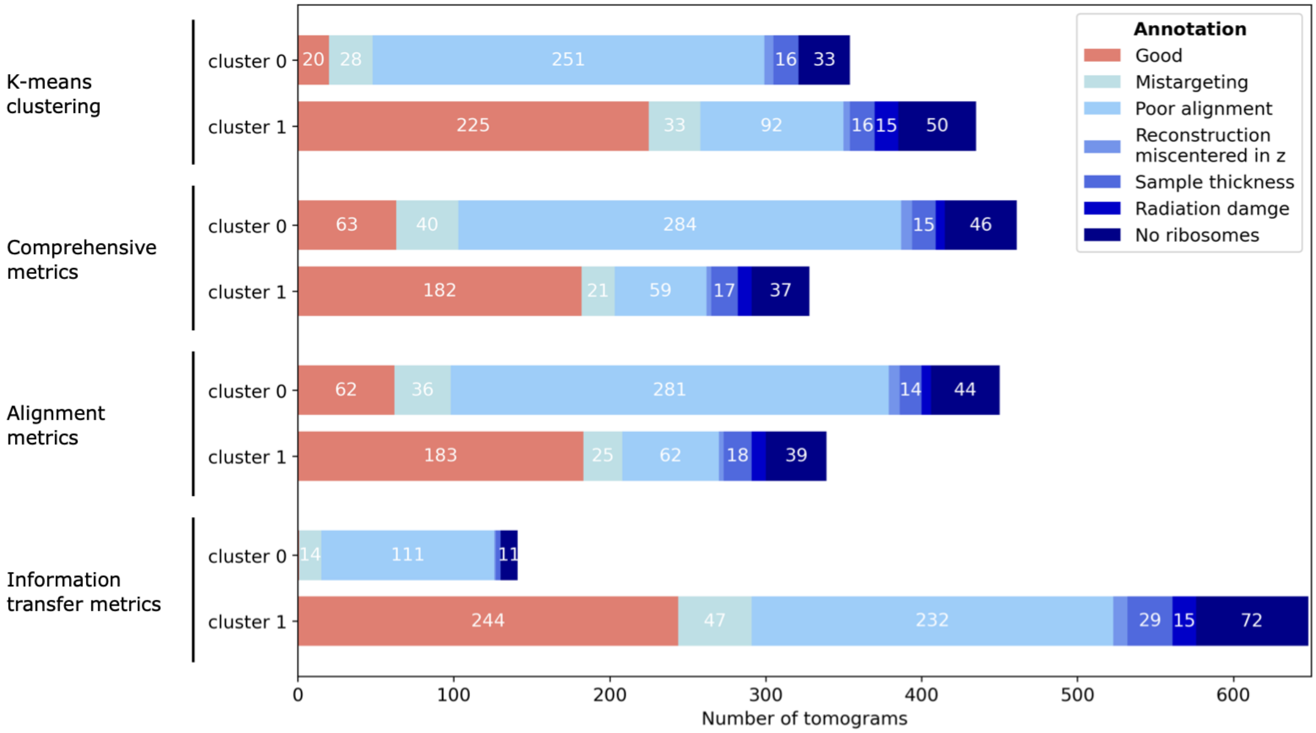


**Extended Data Figure** 10: **Comparative analysis of different approaches for classifying tomograms using AreTomo3's metrics.** Clustering results are compared for different classification strategies for the minicell dataset described in Fig. 4. **a.** $K$-means clustering with 2 clusters was applied, as shown in Fig. 4. **b-d.** Classification was performed based on constant thresholds. The following thresholds were applied as criteria for rejecting tomograms: (i) the estimated tilt axis orientation deviating by more than 0.75° from the dataset's median; (ii) sample thickness exceeding one standard deviation above the median (~360 nm); (iii) maximum global shift exceeding 10% of the tilt series x/y dimensions (~630 Å); (iv) failed local patches exceeding 5% for tilts within ±30° and more than 10% across the full tilt range; and (v) CTF correlation score or (vi) CTF resolution worse than one standard deviation from the median. Classification results are shown based on **b.** all six criteria, **c.** only alignment metrics (i, iii, iv), and **d.** only the information transfer metrics (v, vi).

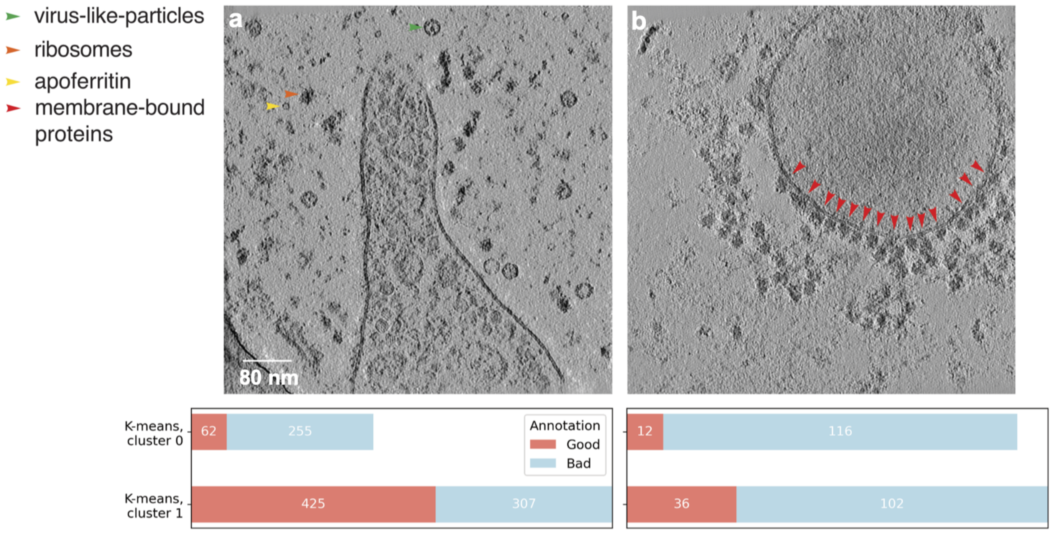


**Extended Data Figure** 11: **Comparison of $k$-means clustering on AreTomo3's metrics and user classification of the phantom**[35] **and affinity-purified lysosome datasets. a.** For the phantom dataset, users selected tomograms that contained sufficient copies of the proteins of interest (as annotated) and appeared well-aligned. 45% (255 of 562) of user-rejected tomograms were captured by the rejected cluster in $k$-means clustering (cluster 0), while 12.7% (62 of 487) of user-selected tomograms were misclassified. As quality metrics were introduced as a feature after visual inspection was performed, the tomograms used for visual inspection and clustering were reconstructed by different version of AreTomo3. **b.** In the affinity-captured lysosome dataset (266 tomograms), users selected tomograms based on alignment quality and the presence of membrane-bound proteins (red arrows). The visual inspection was performed on tomograms reconstructed using Athena (ThermoFisher Scientific). Among the user-rejected tomograms, 53% (116 of 218) were correctly identified by clustering, while 25% (12 out of 48) of user-selected tomograms were misclassified. The higher misclassification rate likely reflects that AreTomo3's metrics do not report on the presence of membrane-bound proteins and that user selection was based on tomograms reconstructed by a different software package.

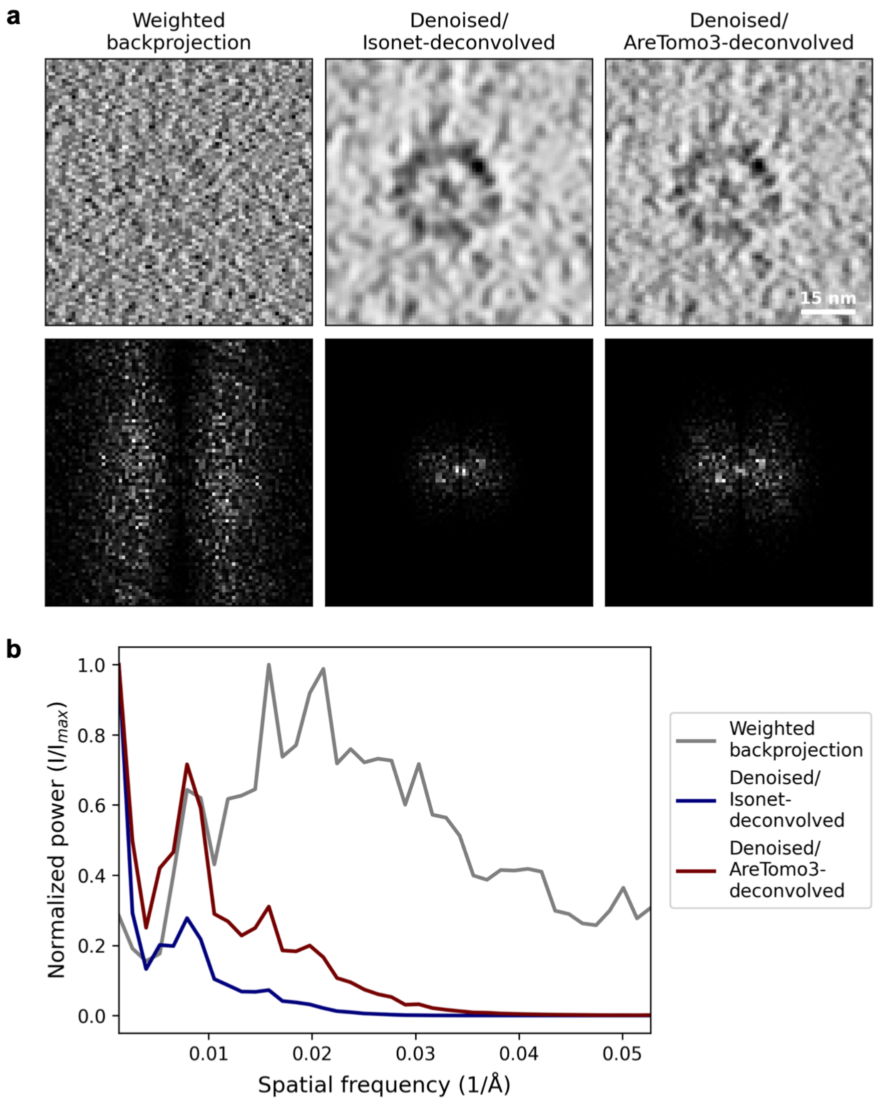


**Extended Data Figure** 12: **AreTomo3's CTF correction enables denoising to retain more intermediate resolution features. a.** A cross-section through a microtubule from a lamella of a cilium and centrosome is shown from the undenoised tomogram (left) and the indicated denoised tomogram, where denoising was performed after CTF deconvolution by either IsoNet (center) or AreTomo3 (right). The lower panel shows the Fourier transform of the cross-section. In real space, the individual protofilaments are better resolved in the AreTomo3 case. Correspondingly in Fourier space, features are observed to higher resolution. **b.** The normalized radial intensity profiles of the microtubules cross-sections show superior retention of intermediate resolution features in the AreTomo3 case compared to IsoNet, which coincide with features in the weighted backprojection profile.

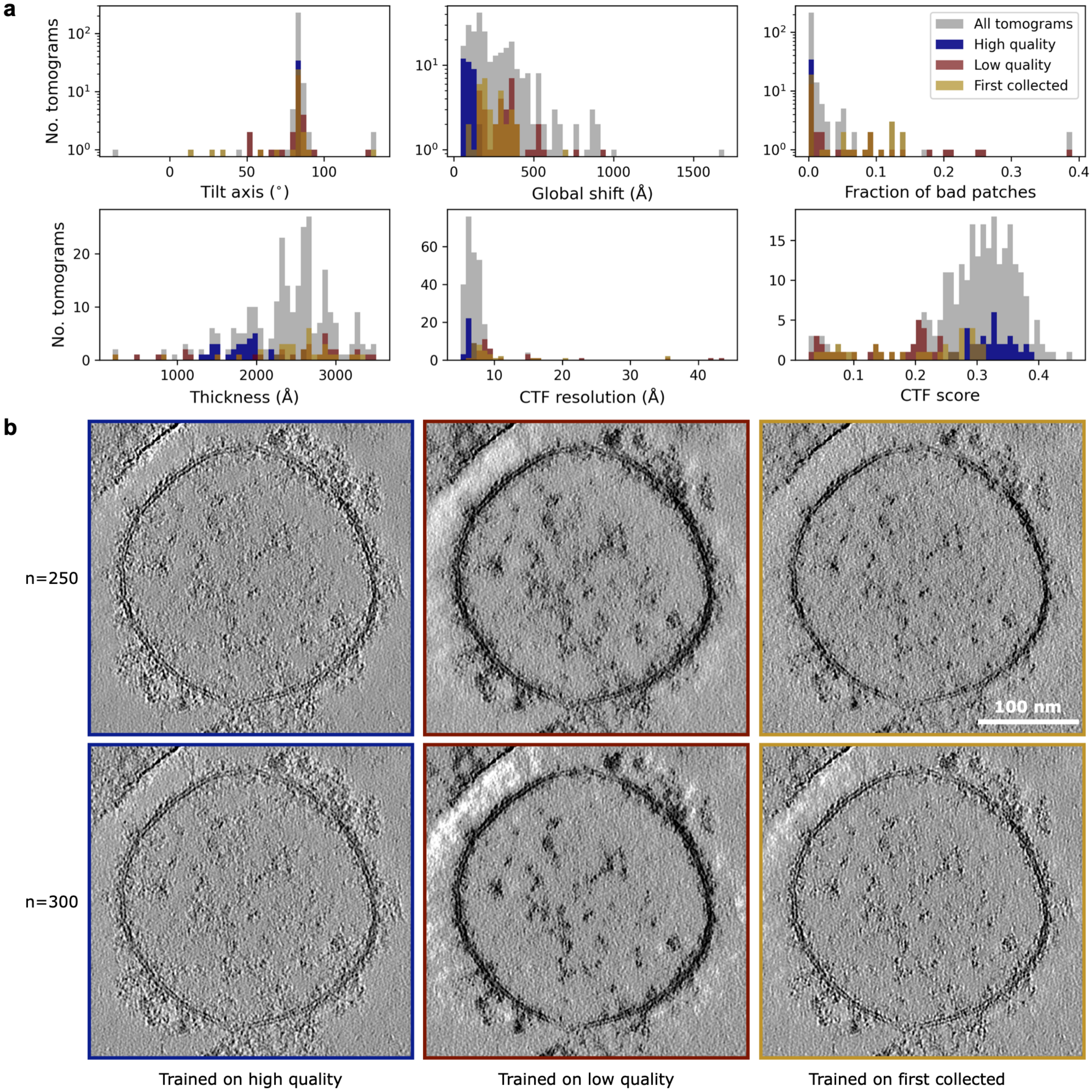


**Extended Data Figure** 13: **Curating training data using AreTomo3's quality metrics yields more robust denoising. a.** Tomograms from a dataset of purified lysosomes were selected if they met certain thresholds for all six quality metrics, resulting in a high quality training set of 34 tomograms (blue). For comparison, training sets were also generated from 34 tomograms with low CTF scores and poor alignment (red) and the first 34 tomograms collected during this session (yellow) to mimic the scenario of running denoising live, except without any data curation. **b.** The resulting models from these different training sets were used to denoise a tomogram that was not included in any of the training sets. To examine consistency across hyperparameter space, the quality of denoising was compared by varying the number of subvolumes extracted per tomogram per training epoch (n) during model training.

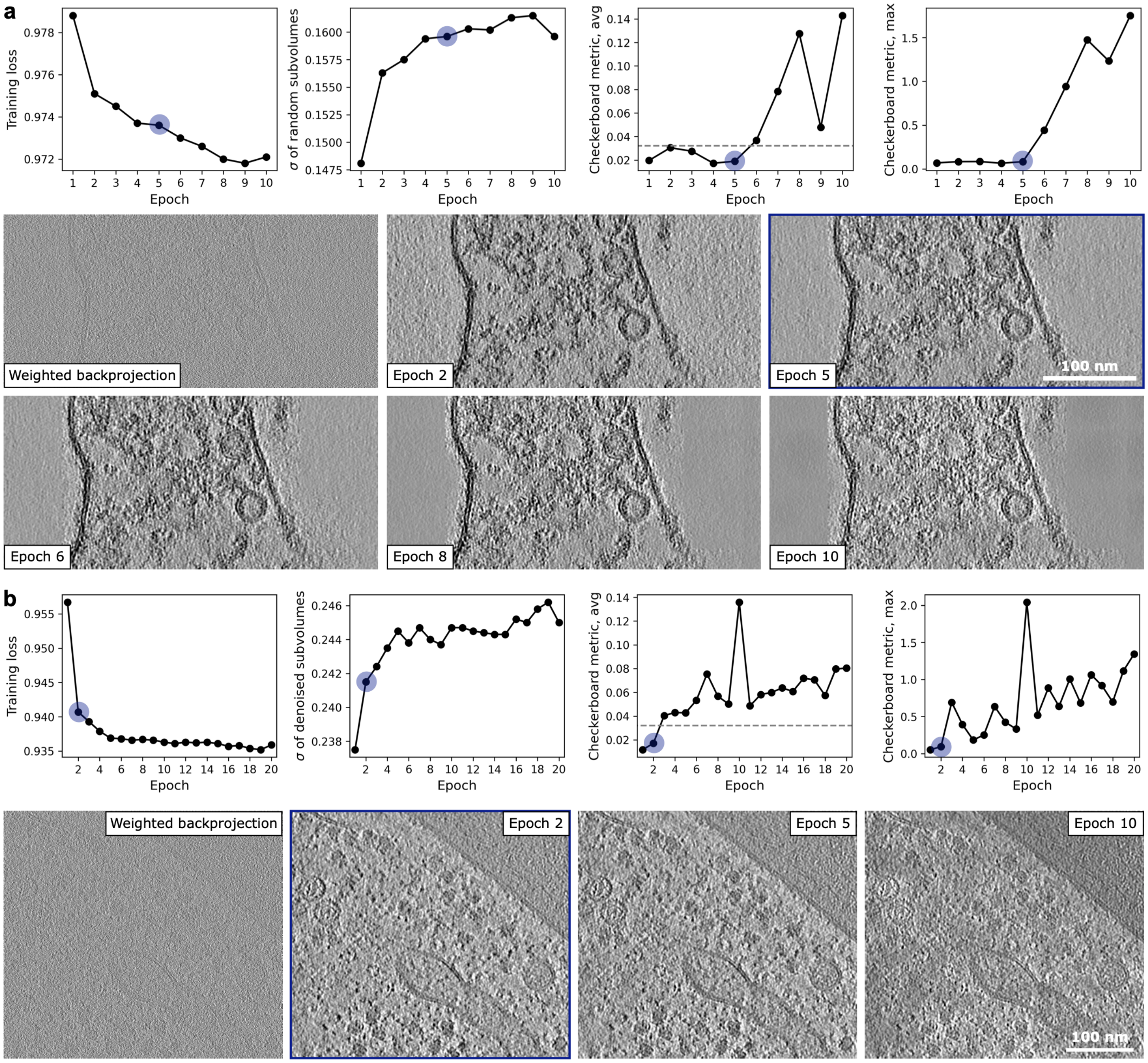


**Extended Data Figure** 14: **Automated termination of training based on the emergence of checkerboard artifacts.** Compared to the training loss and standard deviation of subvolumes from the denoised tomograms, metrics quantifying the appearance of checkerboard artifacts (see Methods) in the denoised tomograms were found to be more sensitive to overdenoising. These metrics are plotted as a function of training epoch (upper), and slices through the indicated tomogram are visualized (lower) for **a.** a dataset of purified synaptosomes and **b.** a dataset of yeast lamella. In both cases, the epoch indicated in blue was chosen based on a maximum threshold of 0.034 for the average checkerboard metric (dashed line).

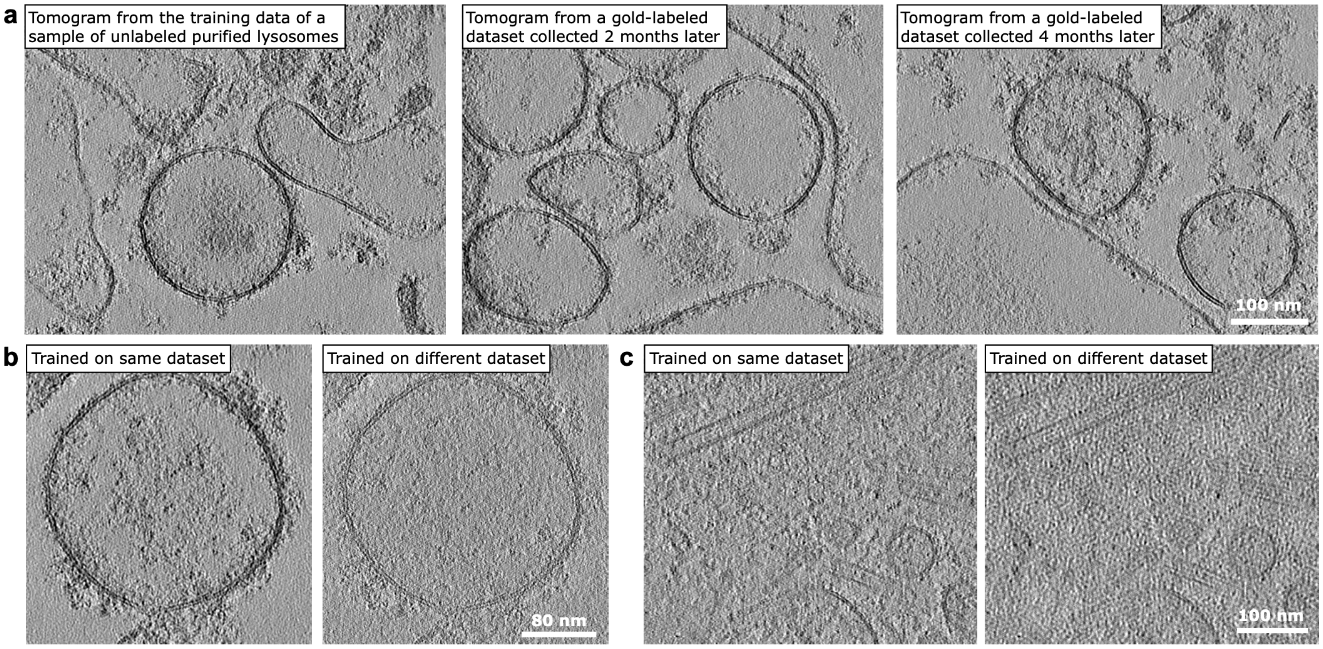


**Extended Data Figure** 15: **Generalizability of denoising models. a.** A model trained on tomograms of affinity-captured unlabeled lysosomes has shown comparable denoising when applied to tomograms from the same data collection session as the training data (left) and to tomograms collected during subsequent data collection sessions from a similar but gold-labeled sample. However, pre-trained models show limited generalizability. **b.** A tomogram of affinity-captured lysosomes was denoised either by this model pre-trained on lysosome data (left) or a model pre-trained on lamella data of a cilium and centrosome (right). In addition to differences in the sample type, the lysosome and cilia data were reconstructed at distinct pixel sizes: 5 and 9.5 Å, respectively. **c.** As a corollary, a tomogram from the cilia dataset was denoised either by the pre-trained cilia model (left) or the pre-trained lysosome model (right).
